# Cdc42 activity is essential for the interplay between cAMP/PKA pathway and CatSper function

**DOI:** 10.1101/2020.08.24.264739

**Authors:** Guillermina M. Luque, Xinran Xu, Ana Romarowski, María G. Gervasi, Gerardo Orta, José L. De la Vega-Beltrán, Cintia Stival, Nicolás Gilio, Tomás Dalotto-Moreno, Dario Krapf, Pablo E. Visconti, Diego Krapf, Alberto Darszon, Mariano G. Buffone

## Abstract

Sperm acquire the ability to fertilize in a process called capacitation and undergo hyperactivation, a change in the motility pattern, which depends on Ca^2+^ transport by CatSper channels. CatSper is essential for fertilization and it is subjected to a complex regulation that is not fully understood. Here, we report that similar to CatSper, Cdc42 distribution in the principal piece is confined to four linear domains and this localization is disrupted in CatSper1-null sperm. Cdc42 inhibition impaired CatSper activity and other Ca^2+^-dependent downstream events resulting in a severe compromise of the sperm fertilizing potential. We also demonstrate that Cdc42 is essential for CatSper function by modulating cAMP production by sAC, providing a new regulatory mechanism for the stimulation of CatSper by the cAMP/PKA-dependent pathway. These results reveal a broad mechanistic insight into the regulation of Ca^2+^ in mammalian sperm, a matter of critical importance in male infertility as well as in contraception.

## Introduction

Sperm acquire the ability to fertilize in the female genital tract in a process called capacitation (Austin, 1952; Chang, 1951). During capacitation sperm sense the environment and undergo a change in the motility pattern called hyperactivation, which is critical for fertilization (Suarez and Ho, 2003). This high-amplitude, asymmetric, whip-like beating pattern of the flagellum enables sperm to migrate through the oviduct and to overcome the oocyte surrounding layers (Demott and Suarez, 1992; Stauss et al., 1995).

Hyperactivation requires a Ca^2+^ uptake through the sperm-specific CatSper channel complex (Ren et al., 2001). This complex comprises four homologous α subunits (CatSper 1–4) (Kirichok and Lishko, 2011; Navarro et al., 2008) and auxiliary subunits: CatSper β, CatSper y, CatSper δ (Chung et al., 2011; Liu et al., 2007; Wang et al., 2009), and the more recently described CatSper ε, CatSper ζ and EFCAB9 (Chung et al., 2017; Hwang et al., 2019). CatSper proteins are encoded by multiple testis-specific genes and localized in the plasma membrane of the principal piece of mature sperm (Lobley et al., 2003; Quill et al., 2001; Ren et al., 2001), except for CatSper ζ and EFCAB9 which are small soluble proteins (Chung et al., 2017; Hwang et al., 2019). Mice lacking any of the CatSper1–4 genes (Qi et al., 2007; Quill et al., 2003; Ren et al., 2001) as well as human males with CatSper loss of function mutations (Avenarius et al., 2009; Avidan et al., 2003; Smith et al., 2013) are infertile and their sperm unable to hyperactivate.

The CatSper channel is probably the most complex of all ion channels characterized at the moment. It provides the fine tuning for Ca^2+^ homeostasis required for changes in motility that sperm experience during their transit to the fertilization site. In addition, CatSper channels are subject to a sophisticated regulation that differs according to the species. For example, exogenous compounds such as progesterone and prostaglandins produce a robust channel activation in human but not mouse sperm (Lishko et al., 2011; Miller et al., 2015) mediated by degradation of lipid endocannabinoid 2-arachidonoylglycerol (2AG) (Miller et al., 2016). In contrast, both mouse and human CatSper channels are activated by intracellular alkalinization (Kirichok et al., 2006; Lishko et al., 2010). This may indicate species-specific adaptations of sperm to adjust to a distinct set of activators within the female reproductive tract (Lishko and Mannowetz, 2018).

Recent reports found that mouse CatSper activity is upregulated by cAMP-dependent activation of protein kinase A (PKA) (Orta et al., 2018). From a molecular point of view, bicarbonate (HCO_3_^-^) and Ca^2+^ stimulation of the soluble adenylyl cyclase (sAC; aka ADCY10) leads to an increase in cAMP, PKA activity and a complex regulation of tyrosine phosphorylation (pY) of sperm proteins (Alvau et al., 2016; Navarrete et al., 2015; Visconti et al., 1995). It remains incompletely understood if PKA itself phosphorylates CatSper or if its activation relays on other intermediary events (Orta et al., 2018). Thus, the interplay between these players persist unresolved.

The complex structural organization of CatSper has impeded the heterologous reconstitution of the channel to study its regulation. By using three-dimensional stochastic optical reconstruction microscopy (3D STORM), it was recently reported that CatSper proteins form a unique pattern of four linear domains in the plasma membrane close to the fibrous sheath, along the principal piece of the flagellum (Chung et al., 2014; Chung et al., 2017; Hwang et al., 2019). This segregated localization of proteins in the tail may provide the structural basis for the selective activation of CatSper and the normal development of hyperactivated motility. For example, recent super-resolution microscopy experiments in human sperm revealed that Hv1, the voltage-gated proton channel involved in cytoplasmic alkalinization, is distributed asymmetrically within bilateral longitudinal lines and that inhibition of this channel leads to a decrease in sperm rotation along the long axis (Miller et al., 2018). In addition, other signaling molecules such as Caveolin-1, phosphorylated Ca^2+^/calmodulin-dependent protein kinase II (P-CaMKII), calcineurin (PPP3C aka PP2B) and EFCAB9 colocalize with CatSper displaying a similar spatial distribution along the principal piece in mouse sperm (Chung et al., 2014; Hwang et al., 2019). Only CatSper has been shown to be critical for this segregated localization since CatSper-null mice do not display the four-column organization of these signaling molecules (Chung et al., 2014). It is anticipated that these proteins form a signaling complex with CatSper that may constitute an important point of regulation for this channel. However, with the exception of EFCAB9 that provides partial pH and Ca^2+^ sensitivity (Hwang et al., 2019), no reports have demonstrated regulatory functions on CatSper activity.

In somatic cells, the Rho-family of small GTPases proteins (Cdc42, Rac, and Rho) govern a variety of important cellular functions including actin cytoskeleton organization (Hall, 2012). The presence of these proteins and some of the downstream effectors has been described in mammalian sperm (Baltiérrez-Hoyos et al., 2012; Delgado-Buenrostro et al., 2005; Ducummon and Berger, 2006; Romarowski et al., 2015) but their role has not been clearly elucidated since the targeted deletion of these proteins results in embryonic lethality (Chen et al., 2000). In this report, we found an unexpected new role for one of the best characterized members of this family: Cdc42. We provide evidence that Cdc42 is a new regulatory protein that localizes in the CatSper complex and has a critical function on its activity.

## Results

### Cdc42 localizes in four longitudinal lines along the flagellum resembling CatSper distribution pattern

The presence of Cdc42 protein in mouse sperm was analyzed by immunoblotting using a specific antibody against this small GTPase. As shown in Fig. 1A, a single band of the expected size for Cdc42 (21.3 kDa) was observed in these cells. Immunofluorescence studies revealed the localization of Cdc42 mainly along the flagellum as well as in the acrosome of mouse sperm (Fig. 1B).

**Figure 1:**
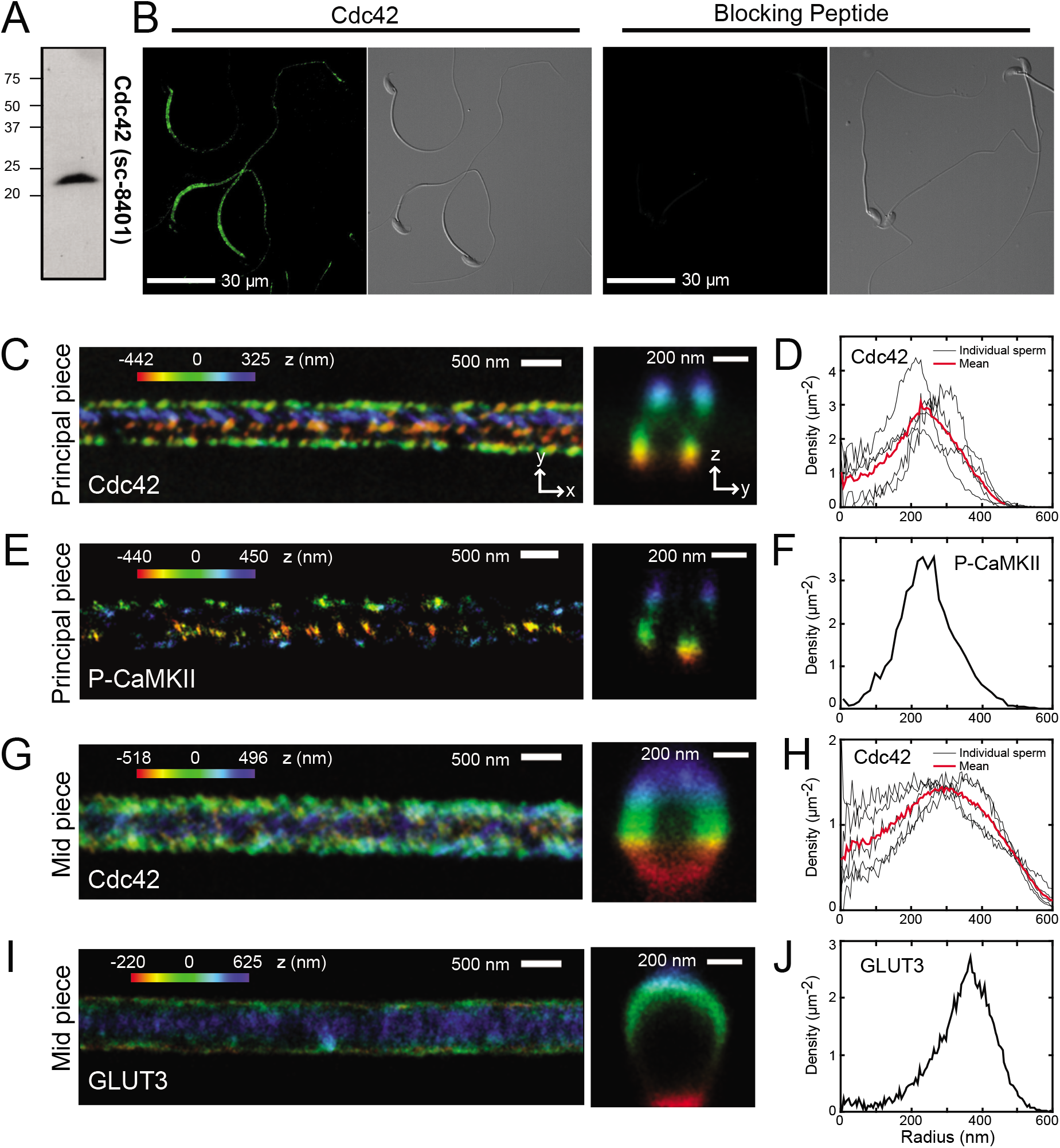
Cdc42 is localized in four longitudinal lines similar to those of the CatSper distribution. **A)** Proteins (equivalent to 14 x 10^6^ sperm) were analyzed by 15% SDS-PAGE and immunoblotted using an antibody against Cdc42 (sc-8401). **B)** Representative immunofluorescence images and their corresponding phase-contrast micrographs of non-capacitated sperm stained with anti-Cdc42 antibody and negative control are shown. Nonspecific staining was determined by incubating the sperm with blocking peptide (sc-8401 P). **C)** Representative 3D STORM images showing the localization of Cdc42 in the sperm principal piece at x-y projection (left) and in y-z cross-section (right). **D)** Radial profiles of Cdc42 localization in principal piece (n=4 sperm). **E)** Representative 3D STORM images of principal piece P-CaMKII. Left, x-y projections. Right, y-z cross-sections. **F)** Radial profiles of P-CaMKII localization in principal piece. **G)** Representative 3D STORM images showing the localization of Cdc42 in the sperm mid piece at x-y projection (left) and in y-z cross-section (right). **H)** Radial profiles of immunostained density of the sperm mid piece localized Cdc42. **I)** Representative 3D STORM images of mid piece GLUT3. Left, x-y projections. Right, y-z cross-sections. **J)** Radial profiles of immunostained density of GLUT3. The color in all x-y projections encode the relative distance from the focal plane along the z axis.

To investigate the spatial distribution of Cdc42, 3D STORM was used. This type of microscopy allows 3D reconstruction with ~20 nm resolution (Huang et al., 2008; Rust et al., 2006), making it possible to study protein localization within the sperm flagellum, which is less than 1 μm in diameter (Chung et al., 2014; Gervasi et al., 2018). The identification of specific structures within the flagellum was performed by analyzing the localization of well-known flagellar proteins. For the axoneme, the fibrous sheath and the plasma membrane, β-tubulin, A kinase anchoring protein 4 (AKAP4), and the facilitative glucose transporter 3 (GLUT3) were used as previously described (Chung et al., 2014) (Suppl. Fig. 1 and 2). Taking advantage of the cylindrical symmetry of the flagellum, the molecule localizations were converted into cylindrical coordinates (*r*, θ, z’) where z’ represents the flagellum axis. In cartesian coordinates (x, y, z), z is the direction normal to the coverslip, so that in this coordinate system the sperm lies on the x,y plane as represented in Suppl. Fig. 1A. As expected, the spatial distribution of β-tubulin was axonemal and localized in the center of the flagellum, while AKAP4 (localized in the fibrous sheath) and GLUT3 (present throughout the flagellar plasma membrane) showed a continuous rim distribution in cross-section (Suppl. Fig. 1B, right panel). The radial distributions of β-tubulin, AKAP4 and GLUT3 in the principal piece showed that β-tubulin peaked at 23 ± 2 nm (mean ± standard error) from the center of the flagellum while AKAP4 and GLUT3 peaked at 146 ± 1 nm and 269 ± 1 nm, respectively (Suppl. Fig. 1C).

**Figure 2:**
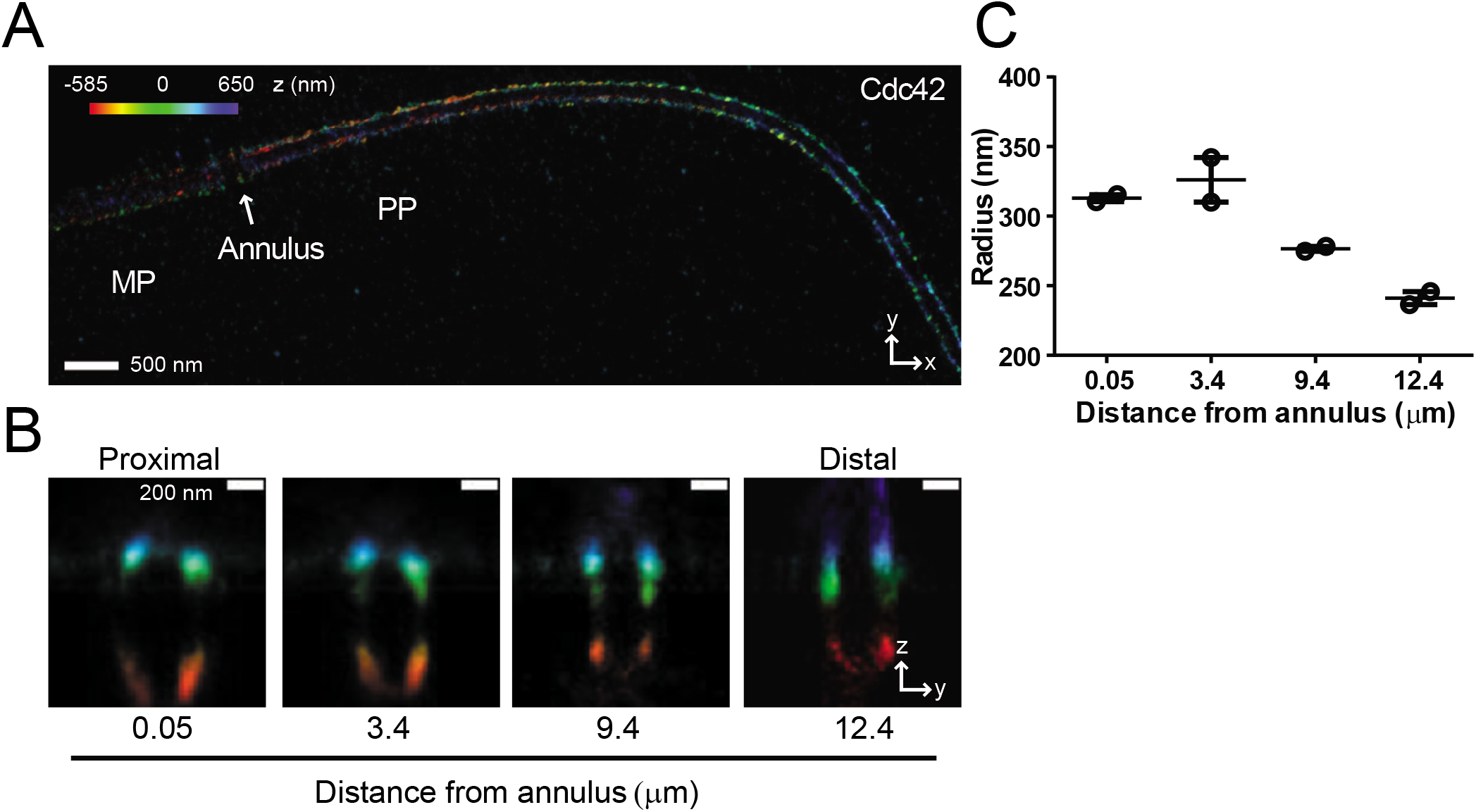
Cdc42 is localized in four longitudinal lines along the principal piece. **A)** Representative 3D STORM images showing the localization of Cdc42 along the sperm principal piece. Entire x-y field of view. MP, mid piece. PP, principal piece. An arrow marks the annulus. The color encode the relative distance from the focal plane along the z axis. **B)** Series of magnified y-z cross-sections obtained from the proximal to the distal part of the principal piece (same cell). Distance from annulus (μm) is specified. **C)** The radius (nm) of Cdc42 staining was calculated in the principal piece at different distances from the annulus.. Values represent the mean ± SEM of 2 measurements.

Mouse sperm exhibited a Cdc42 localization confined to four columns along the principal piece (Fig. 1C and Suppl. Video 1), which resembles the localization of CatSper in the same region (Chung et al., 2014). On cross-sections of the flagellum, Cdc42 appeared as four tight puncta (Fig. 1C right panel). Similar to Cdc42, P-CaMKII showed the CatSper-like distribution in the principal piece (Fig. 1E-F and Suppl. Fig. 3C-D), as previously described (Chung et al., 2014). The radial distributions of Cdc42 and P-CaMKII peaked at 238 ± 16 nm and 235 ± 2 nm respectively (Fig. 1D and F). These values are similar to those observed for GLUT3 (Suppl. Fig. 1B-C), indicating that both proteins localized close to the plasma membrane. A graphical representation of Cdc42 localization in terms of the azimuth angle θ vs. axial distance z’ along the principal piece showed the four stripes along the flagellum (Suppl. Fig. 3A) as well as the 2D angular profiles of the surface localizations (Suppl. Fig. 3B). This segregated localization of Cdc42 in the principal piece, disappeared in the flagellum mid piece, where a continuous ring staining was observed in the cross-section (Fig. 1G). The radial distribution in this section peaked at 281 ± 25 nm (Fig. 1H), while GLUT3 was observed at 323 ± 2 nm in that region (Fig. 1I-J).

Since the sperm flagellum is usually close to 1 μm wide in the proximal portion and tapers gradually towards the distal tip, the molecular distribution of Cdc42 across the entire flagellum was visualized with 3D STORM. The localization of Cdc42 in four tight clusters was conserved along the principal piece (Fig. 2). A series of cross-sections was analyzed (Fig. 2B), where the decrease in Cdc42 radial distribution as the distance from the annulus increases was observed (Fig. 2C).

Altogether, these results indicate that Cdc42 is organized in four columns in the principal piece and resembles the localization of CatSper and other signaling molecules reported in that structural domain.

### Cdc42 is delocalized in CatSper1 KO sperm

It has been shown that CatSper channel complex serves as an organizing structure for the four-line domains. In this respect, other molecules such as Caveolin-1, P-CaMKII and PP2B lost their localization when CatSper is abrogated (Chung et al., 2014). Therefore, to test the extent by which Cdc42 is associated to this complex, its localization in sperm from CatSper1-null mice was analyzed. Levels of protein expression in CatSper1 KO sperm were examined by immunoblotting using two specific antibodies (Fig. 3A-B). Expression of Cdc42 was detected in CatSper1 KO, albeit at 50% lower levels than in HET sperm (Fig. 3A-B, right panel). Immunofluorescence studies revealed that Cdc42 was also present along the flagellum and in the acrosome of CatSper1 KO (Fig. 3C) as in the HET sperm; although with lower fluorescence intensity, consistent with reduced protein expression (Fig. 3A-B). 3D STORM analyses in principal piece from CatSper1 KO sperm were performed, but only poor-quality reconstructions could be obtained due to the low number of localized fluorophores (2 ± 1 localizations/nm in CatSper1 KO vs. 10 ± 2 localizations/nm in HET sperm), in agreement with lower protein expression. In summary, 21 out of 23 sperm analyzed of CatSper1 KO sperm showed a clear Cdc42 delocalization from the quadrilateral structure in comparison with HET sperm (Fig. 3D). All together, these results suggest that the absence of CatSper channels alters the expression levels and the spatial 3D distribution of Cdc42.

**Figure 3:**
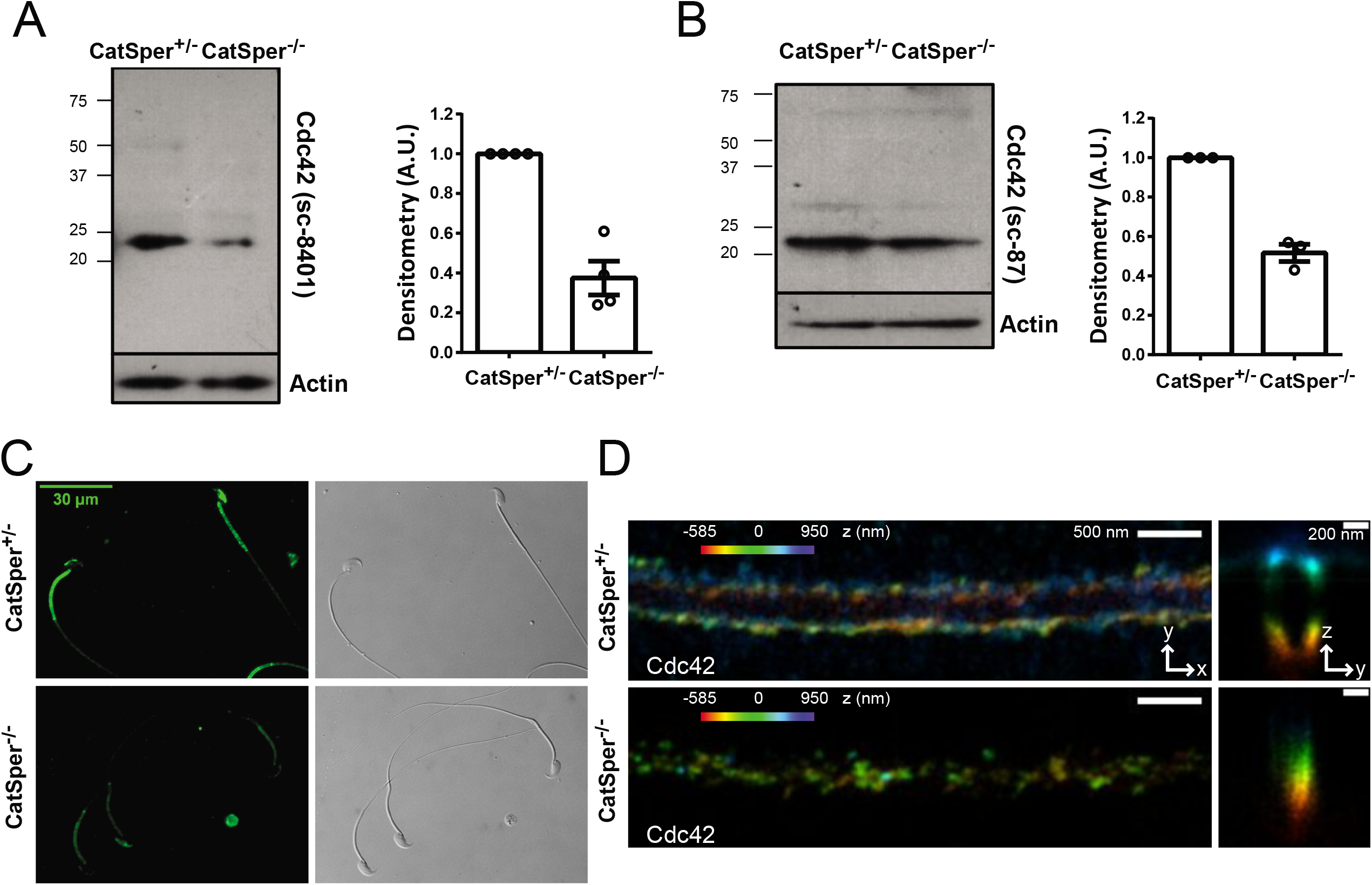
CatSper is essential for Cdc42 expression and spatial organization. **A)** Proteins (equivalent to 12 x 10^6^ sperm) were analyzed by 15% SDS-PAGE and immunoblotted using antibody against Cdc42 (sc-8401). As a loading control, an anti-Actin antibody was used. Quantitative analysis was performed by measuring the optical density of all bands and relativized to Actin. Values represent the mean ± SEM of 4 independent experiments, where normalization to the control condition (CatSper1^+/-^ sperm) was used. Non-parametric Wilcoxon signed-rank test was performed (hypothetical value: 1; control condition). **B)** Proteins (equivalent to 10 x 10^6^ sperm) were analyzed by 15% SDS-PAGE and immunoblotted using antibody against Cdc42 (sc-87). As a loading control, an anti-Actin antibody was used and the quantitative analysis performed as described above. Values represent the mean ± SEM of 3 independent experiments, where normalization to the control condition (CatSper1^+/-^ sperm) was used. Non-parametric Wilcoxon signed-rank test was performed (hypothetical value: 1; control condition). **C)** Representative immunofluorescence image and the corresponding phase-contrast micrographs of non-capacitated control (CatSper1^+/-^) and CatSper1 KO (CatSper1^-/-^) sperm stained with anti-Cdc42 antibody (sc-8401) are shown. **D)** Representative 3D STORM images showing the localization of Cdc42 in the principal piece of control (CatSper1^+/-^) sperm at x-y projection (left, upper panel) and in y-z cross-section (right, upper panel). Representative 3D STORM images of CatSper1 KO (CatSper1 ^-/-^) principal piece Cdc42 (lower panel). Left, x-y projections. Right, y-z cross-sections (n=23 sperm from 3 different mice). The color in all x-y projections encode the relative distance from the focal plane along the z axis.

### Inhibition of Cdc42 decreases [Ca^2+^]_i_ during capacitation

Given the Cdc42 distribution in four linear domains in the principal piece (Fig. 1C-D), we postulated that this signaling protein could be involved in CatSper function. This hypothesis was tested by different approaches using a pharmacological inhibitor. Cdc42 activity is highly regulated by switching between its active GTP-bound form and its inactive GDP-bound form. This modulation depends on the exchange of GDP to GTP by guanine nucleotide exchange factors (GEF), whereas GTPase activating proteins (GAP) bind to active Cdc42 stimulating the GTP-hydrolytic reaction, thus terminating the signaling event (Rossman et al., 2005). In addition, guanine nucleotide dissociation inhibitors (GDI) prevent GEF-mediated nucleotide exchange, maintaining the GTPase in an inactive state and promoting Cdc42 dissociation from the membrane (Garcia-Mata et al., 2011). To assess the function of Cdc42 in mouse sperm, MLS-573151, a specific pharmacological inhibitor that prevents the binding of GTP to Cdc42 (Surviladze et al., 2010) was used.

To analyze the participation of Cdc42 on CatSper activity, [Ca^2+^]_i_ was evaluated by flow cytometry in the presence or absence of MLS-573151. We recently determined that the capacitation-associated [Ca^2+^]_i_ increase at 90 min was dependent of CatSper channels, as sperm derived from CatSper1 KO failed to increase [Ca^2+^]_i_ (Luque et al., 2018). Thus, [Ca^2+^]_i_ was measured by flow cytometry in live sperm loaded with the Ca^2+^ sensitive probe Fluo-4 AM. Sperm were incubated for 90 min under capacitating conditions in the presence of increasing concentrations of MLS-573151. Addition of MLS-573151 significantly suppressed the rise in [Ca^2+^]_i_ that occurs during capacitation as a result of CatSper activation (Fig. 4A-B) in a concentration dependent manner (Fig. 4C and Suppl. Fig. 4). The concentration of 20 μM of MLS-573151 was chosen for all the experiments as the [Ca^2+^]_i_ blockage was maximum without changing sperm viability. This inhibition can be visualized by a decrease in the normalized median fluorescence intensity (MFI) of Fluo-4 (Suppl. Fig. 4) as well as in the percentage of sperm that displayed a rise in [Ca^2+^]_i_ (Fig. 4C). The population of sperm with high fluorescence was determined from the capacitating control condition (CAP 90 min) and extrapolated to the other conditions of each experiment (Fig. 4B).

**Figure 4:**
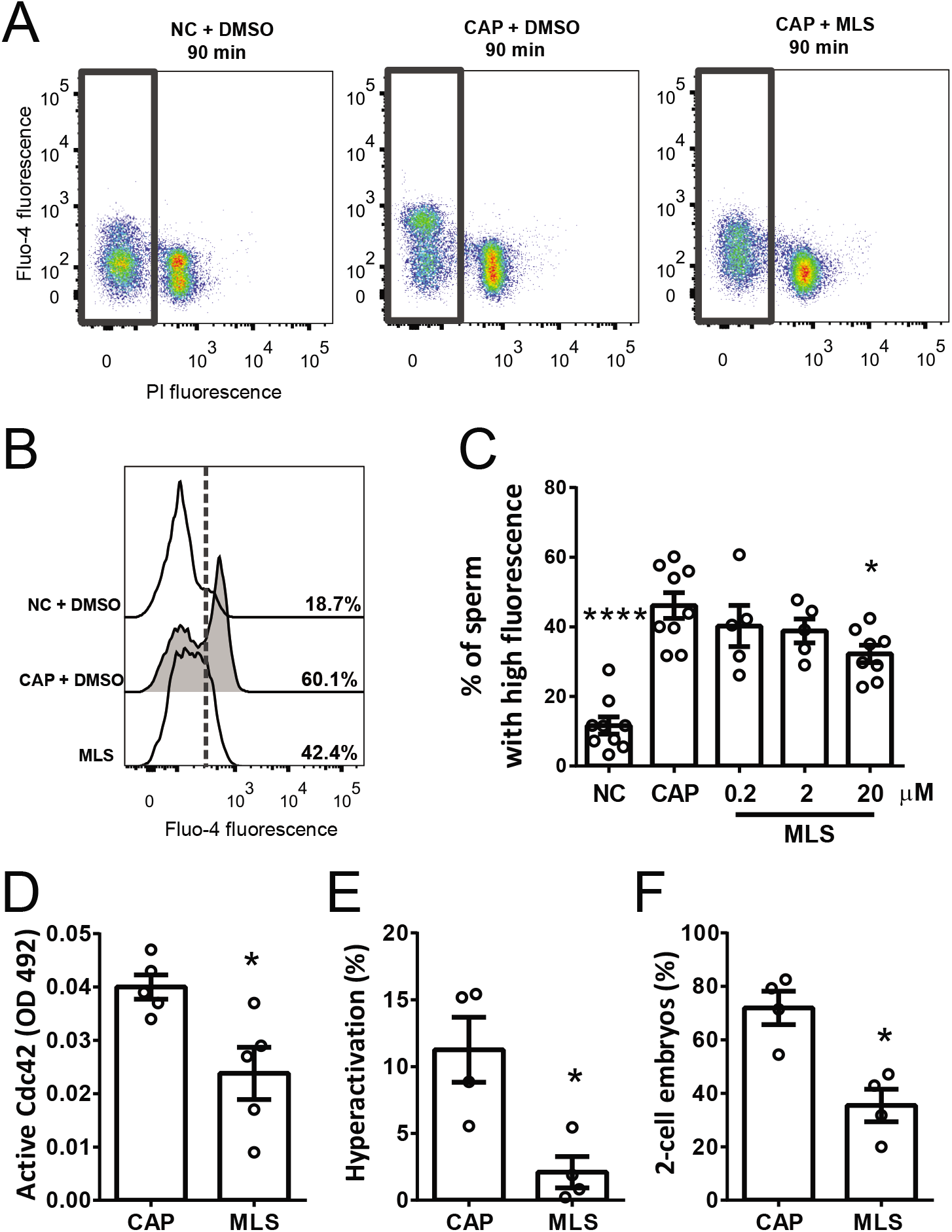
Capacitation associated [Ca^2+^]_i_ increase, hyperactivation and *in vitro* fertilization were block by Cdc42 inhibitor. Sperm incubated for 90 min under non-capacitating (NC) or capacitating conditions (CAP) in the absence (DMSO) or presence of MLS-573151 were analyzed. **A)** Representative two-dimensional Fluo-4 vs. PI fluorescence dot plot are shown. Live sperm (with low PI fluorescence levels) were identified (delimited by gray rectangle) from which Fluo-4 fluorescence levels were analyzed in order to stablished [Ca^2+^]_i_. A concentration of 20 μM of MLS-573151 was used. **B)** Representative histograms of normalized frequency vs. Fluo-4 fluorescence of non-PI stained sperm (live), with the corresponding percentage of sperm that increased Fluo-4 fluorescence, are shown. The percentage of sperm that responds by increasing the [Ca^2+^]_i_ was established in the CAP control condition and extrapolated to the others conditions (dashed line). A concentration of 20 μM of MLS-573151 was used. **C)** Percentage of sperm that increased Fluo-4 fluorescence after being incubated for 90 min in the different conditions. MLS-573151 produce a concentration-dependent decrease in [Ca^2+^]_i_. Values represent the mean ± SEM of 9 independent experiments. **** p<0.0001, * p<0.05 represents statistical significance vs. control (CAP 90 min with DMSO). One-way ANOVA with Dunnett’s multiple comparisons test was performed. **D)** Quantification of active (GTP-bound) Cdc42 in the presence or absence of 20 μM MLS-573151. Values represent the mean ± SEM of 5 independent experiments. * p<0.05 represents statistical significance vs. control (CAP 90 min with DMSO). Paired t-test was performed. **E)** Percentage of hyperactive sperm (%). A concentration of 20 μM of MLS-573151 was used. Data represents the mean ± SEM of 4 independent experiments. * p<0.05 represents statistical significance vs. control (CAP 90 min with DMSO). Unpaired t-test was performed. **F)** Fertilization rates after *in vitro* fertilization (percentage of 2-cell embryos per total oocytes examined). A concentration of 20 μM of MLS-573151 was used. Data represents the mean ± SEM of at least 4 independent experiments. * p<0.05 represents statistical significance vs. control (CAP 90 min with DMSO). Unpaired t-test was performed.

To corroborate that MLS-573151 is inhibiting Cdc42 activity, an ELISA-based assay which measure the GTP-bound form of Cdc42 was performed. As expected, sperm capacitated in the presence of MLS-573151 significantly decreased the Cdc42 active form (Fig. 4D).

### Cdc42 activity is required for hyperactivation and in vitro fertilization

Ca^2+^ brought in from CatSper channels is essential for hyperactivation and successful fertilization (Ren et al., 2001). Given the decrease in [Ca^2+^]_i_ observed as a result of Cdc42 inhibition, hyperactivated motility and *in vitro* fertilization was assessed in presence of MLS-57151. For this purpose, motility patterns were analyzed by computer-assisted semen analysis (CASA) in those samples exposed to Cdc42 inhibitor or control capacitating medium. Cdc42 inhibition by MLS-573151 produced a significant decrease in the percentage of hyperactivated cells when compared with controls (Fig. 4E). This decrease was also observed in the kinetic parameters evaluated, in particular, in those that are used to identify the hyperactivated population such as VCL, LIN and ALH (Suppl. Table 1). Although a slight decrease in sperm motility was observed with the different treatments (Suppl. Table 1), the high percentage of motile sperm observed after exposure to Cdc42 inhibitor (more than 80%) indicated that sperm viability was not affected by this compound.

Next, *in vitro* fertilization of cumulus oocyte complexes (COC) was performed using sperm that were capacitated in the presence or absence of Cdc42 inhibitor, and the percentage of fertilized eggs was determined. Fertilization rates were significantly lower when Cdc42 was inhibited by MLS-573151, supporting its role on capacitation (Fig. 4F).

### Cdc42 activity is essential for CatSper function

To determine whether Cdc42 activity is required for CatSper function, we used a fluorescence method as previously described (Ernesto et al., 2015). This assay is based on the fact that removing external Ca^2+^ and Mg^2+^ after adding EGTA allows CatSper to efficiently conduct monovalent cations (Kirichok et al., 2006), where a sudden influx of Na^+^ depolarizes the cells (Espinosa and Darszon, 1995; Torres-Flores et al., 2011). The magnitude of this depolarization mainly depends on the extent of CatSper opening, and is inhibited by CatSper channels blockers (Ernesto et al., 2015; Torres-Flores et al., 2011). Sperm membrane potential (Em) was measured with a fluorescent cyanine dye DiSC_3_(5). Because this cationic dye accumulates on cells with hyperpolarized membranes, an increase in DiSC_3_(5) fluorescence in the medium reflects Em depolarization. Thus, after EGTA addition, the magnitude of the Em depolarization was visualized as changes in DiSC_3_(5) fluorescence in the extracellular medium. Incubation with Cdc42 inhibitor during capacitation significantly diminished the Em depolarization caused by EGTA addition (Fig. 5A-B) suggesting that CatSper opening is affected by inhibiting Cdc42.

**Figure 5:**
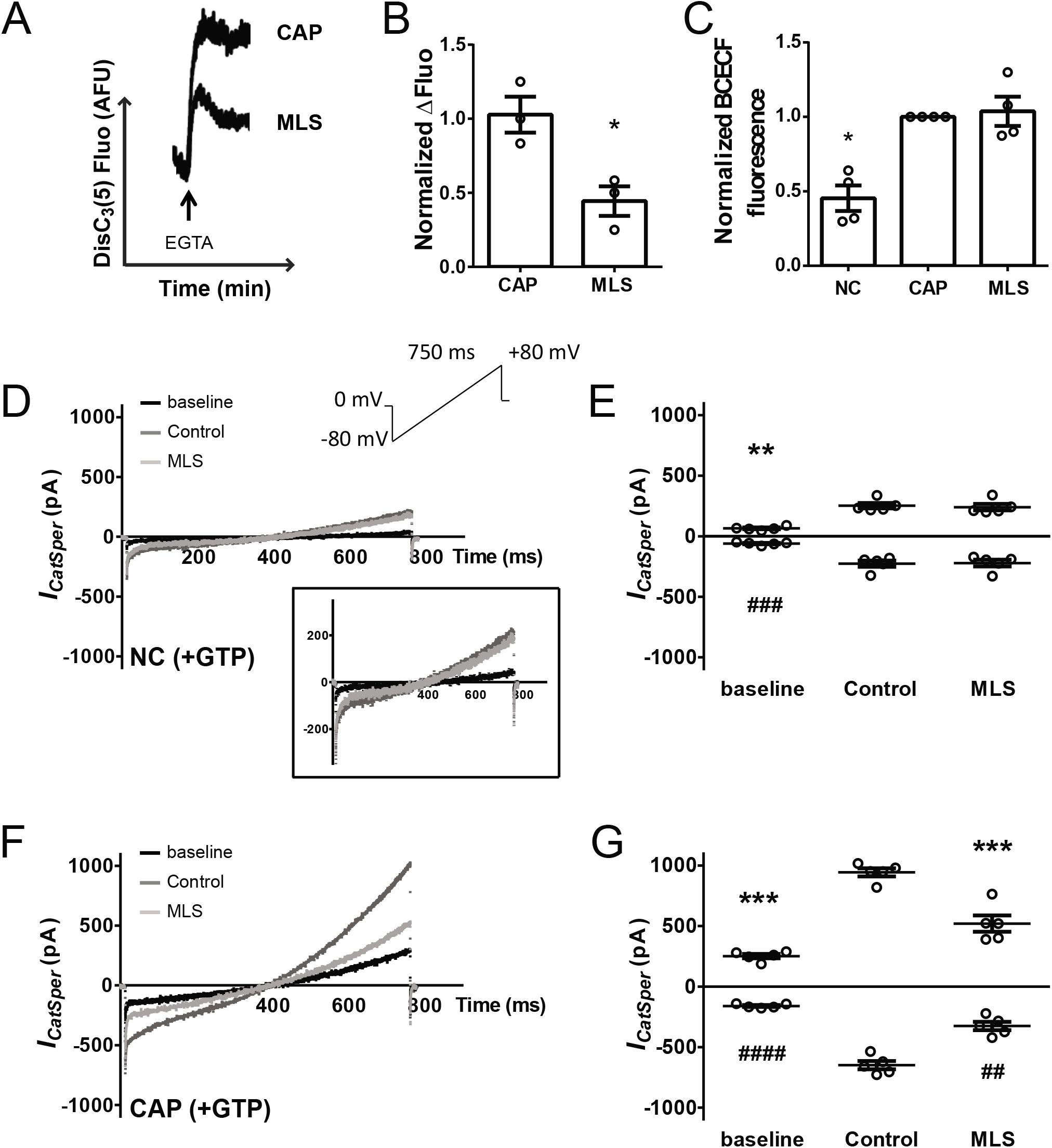
Capacitation associated increase in CatSper currents was block by Cdc42 inhibitor. **A)** Sperm were incubated for 90 min under capacitating conditions (CAP) in the absence (DMSO) or presence of Cdc42 inhibitor (10 μM MLS-573151). CatSper activity was analyzed by measuring Em of sperm loaded with DiSC_3_(5), before and after addition of 3.5 mM EGTA. Representative DiSC_3_(5) fluorescence traces (AFU: arbitrary fluorescence units) through time are shown. The depolarization caused by addition of EGTA is diminished by MLS-573151. **B)** Normalized △Fluorescence represents the difference between DiSC_3_(5) fluorescence after EGTA addition (F_EGTA_) and before (resting: F_R_) compared to the mean obtained in the control condition (CAP 90 min with DMSO). Data represent the mean ± SEM of 3 independent experiments. * p<0.05 represents statistical significance vs. control (CAP 90 min with DMSO). Unpaired t-test was performed. **C)** Mouse sperm pH_i_, was measured by using BCECF AM and flow cytometry. Sperm incubated for 90 min under non-capacitating (NC) or capacitating conditions (CAP) in the absence (DMSO) or presence of Cdc42 inhibitor (20 μM MLS-573151) were analyzed. Normalized MFI of BCECF compared to the control condition (CAP 90 min with DMSO). Values represent the mean ± SEM of 4 independent experiments. * p<0.05 represents statistical significance vs. control (CAP 90 min with DMSO). Non-parametric Kruskal-Wallis test was performed in combination with Dunn’s multiple comparisons test. **D, F)** Representative whole-cell currents traces from non-capacitated (**D**) and capacitated (**F**) mouse sperm. Inward and outward currents were elicited by a voltage ramp protocol from a holding potential of 0 mV (inset). To ensure stable recording conditions, first baseline currents (in HS solution) were obtained. Under HS condition (black traces) CatSper currents were minimal. In cation divalent-free (DVF) medium (dark gray traces) typical CatSper monovalent currents can be recorded. In presence of 20 μM MLS-573151 (light gray traces), CatSper currents were inhibited only under CAP conditions. The patch pipette was always filled with 0.4 mM GTP. **E, G)** Quantification of *I_CatSper_* current densities for all three conditions. MLS-573151 (20 μM) did inhibit the capacitation-activated CatSper current at both negative and positive potentials (−80 and +80 mV). Data represents mean ± SEM of 5 sperm from different mice. *** p<0.001, ** p<0.01 represents statistical significance vs. DVF control (+80 mV). #### p<0.0001, ### p<0.001, ## p<0.01 represents statistical significance vs. DVF control (−80 mV). One-way ANOVA was performed in combination with Holm-Sidak’s multiple comparisons test.

CatSper channels are strongly activated by an intracellular alkalinization (Kirichok et al., 2006; Lishko et al., 2010). To evaluate if Cdc42 alters the sperm intracellular pH (pH_i_), this parameter was analyzed in live cells by flow cytometry, loading sperm with the pH_i_ sensitive probe BCECF AM. Cells were incubated for 90 min under non-capacitating and capacitating conditions in the presence or absence of Cdc42 inhibitor. The intracellular alkalinization-associated with capacitation was not modified by the presence of MLS-573151 (Fig. 5C).

To better characterize the Cdc42-dependent CatSper gating, whole-cell voltage-clamp recordings of CatSper currents (*I_CatSper_*) were performed. *I_CatSper_* from cauda epididymal sperm were analyzed using a typical voltage-ramp protocol (from −80 to +80 mV during 750 ms with a holding potential of 0 mV) and Na^+^ out/Cs^+^ in as the main conducting ions in the absence of divalent cations (divalent cation free condition, DVF). A concentration of 0.4 mM GTP was added to the recording pipette solution to ensure that the activity of Cdc42 is not limited by the lack of this cofactor. Under non-capacitating conditions, the *I_CatSper_* was not altered by the presence of MLS-573151 (Fig. 5D-E). However, while *I_CatSper_* was strongly stimulated under capacitating conditions, this current was potently decreased by the addition of MLS-573151 (Fig. 5F-G). All together, these results suggest that Cdc42 activity is necessary for CatSper function, by a mechanism different from pH_i_ regulation.

There is evidence that CatSper channels are promiscuous and may be inhibited by different compounds in a non-specific manner (Barratt and Publicover, 2012; Brenker et al., 2012). To validate our results, we explored the specificity of the Cdc42 inhibitor used in previous experiments. To this aim, we used a condition where Cdc42 activity is compromised (absence of GTP) and took advantage of the fact that mouse CatSper is strongly activated by intracellular alkalinization (promoted by NH_4_Cl addition), as demonstrated by an increase in *I_CatSper_* (Kirichok et al., 2006). The mechanism for pH-dependent activation is not yet fully understood but it has been reported that the protein EFCAB9 together with the highly rich histidine CatSper amino terminus domain are responsible for sensing changes in pH_i_ (Hwang et al., 2019; Ren et al., 2001). Our experiment is based on the premise that if CatSper is non-specifically blocked by the action of Cdc42 inhibitor, alkalinization would not produce a robust increase in CatSper activity.

In order to analyze whether Cdc42 modulation on CatSper disturbs pH_i_ activation, non-capacitated sperm loaded with the Em sensitive probe DiSC_3_(5) were first exposed to Cdc42 inhibitor and then to NH_4_Cl. CatSper opening was evidenced by the magnitude of Em depolarization resulting from Ca^2+^ influx. MLS-573151 pre-incubation did not alter CatSper sensitivity to pH alkalinization (Fig. 6A-B).

**Figure 6:**
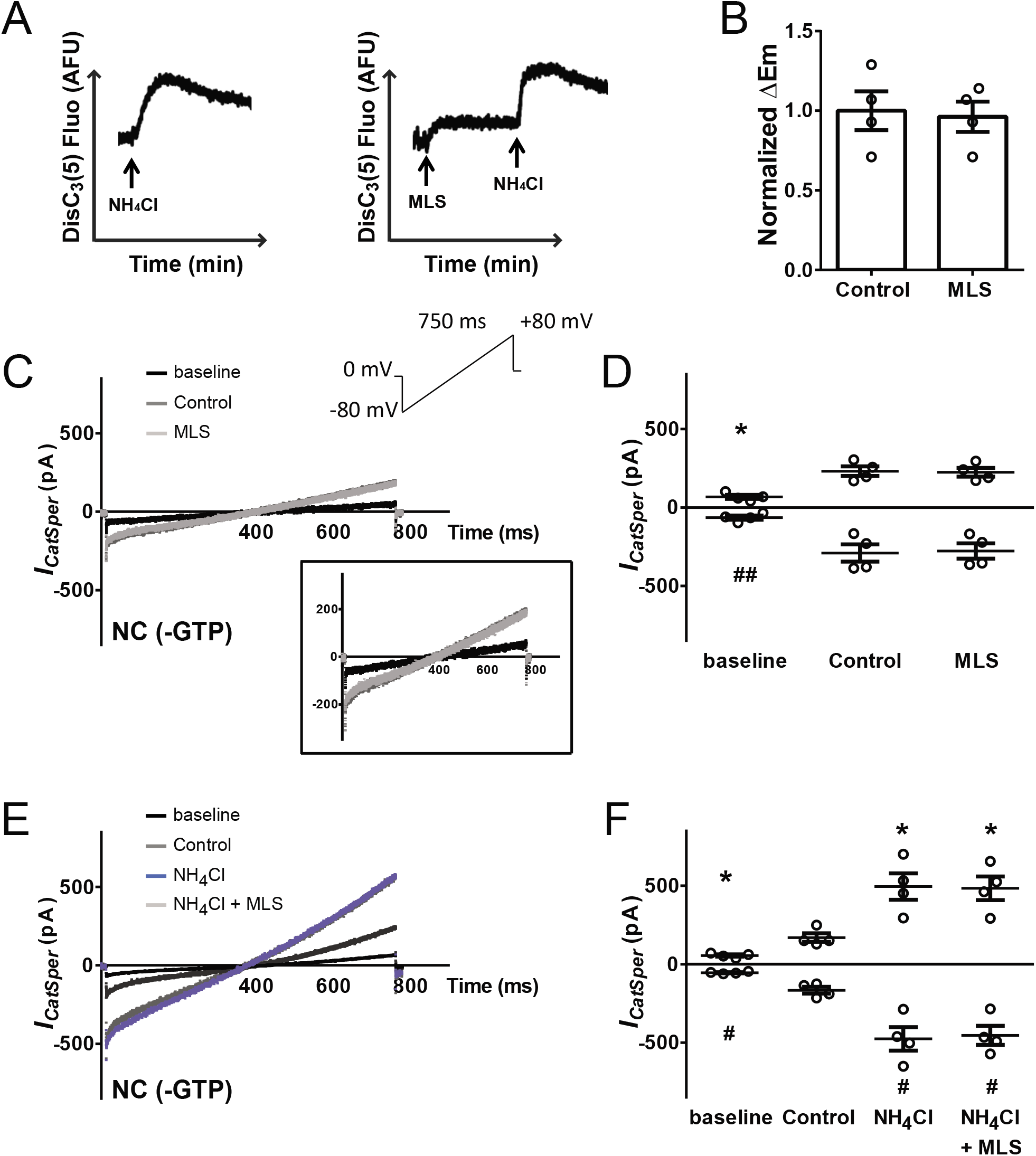
Alkalinization-activated CatSper current is not modified by MLS-573151. CatSper activity was analyzed by measuring Em of sperm obtained from swim-out with DiSC_3_(5) in a population assay. **A)** Representative fluorescence traces (AFU: arbitrary fluorescence units) of Em through time are shown. The addition of 10 mM NH_4_Cl activates CatSper channel causing depolarization by Ca^2+^ influx (left traces). Addition of 10 μM MLS-573151 previous to NH_4_Cl 10 mM did not modified CatSper activation (right traces). **B)** Normalized △Em represents the difference between Em after NH_4_Cl addition (Em_NH4CI_) and before (resting: EmR or Emcdc42 inhibitor) compared to the mean obtained in the control condition (without Cdc42 inhibitor addition). Data represents the mean ± SEM of at least 3 independent experiments. △Em of NC = 14.91 ± 0.96 mV (n = 11). Unpaired t-test was performed. **C, E)** Representative whole-cell currents traces from non-capacitated mouse sperm without addition of GTP to the recording pipette. Inward and outward currents were elicited by a voltage ramp protocol from a holding potential of 0 mV (inset). To ensure stable recording conditions, first baseline currents (in HS solution) were obtained. Under HS condition (black traces) CatSper currents were minimal. In cation divalent-free (DVF) medium (dark gray traces) typical CatSper monovalent currents can be recorded. **C)** In presence of 20 μM MLS-573151 (light gray traces), CatSper currents were not modified. **E)** DVF after adding 10 mM NH_4_Cl, with (light gray traces) or without (blue traces) 20 μM MLS-573151. **D, F)** Quantification of *I_CatSper_* current densities. MLS-573151 (20 μM) did not alter the NH_4_Cl stimulated *I_CatSper_*. Data represents mean ± SEM of 4 sperm from different mice. * p<0.05 represents statistical significance vs. DVF control (+80 mV). ## p<0.01, # p<0.05 represents statistical significance vs. DVF control (−80 mV). One-way ANOVA was performed in combination with Holm-Sidak’s multiple comparisons test.

To further corroborate this result, *I_CatSper_* was analyzed under non-capacitating conditions, without addition of GTP in the recording patch-clamp pipette solution, where the *I_CatSper_* was not altered by MLS-573151 (Fig. 6C-D). The increase in *I_CatSper_* as a result of CatSper activation induced by pH_i_, alkalinization was obtained by adding NH_4_Cl (10 mM), and this stimulation was not modified by the presence of MLS-573151 (Fig. 6E-F). Overall, MLS-573151 did not directly affect *I_CatSper_* nor the alkalinization induced Em depolarization under these experimental conditions. These results support that this inhibitor is not producing a non-specific blockade of CatSper channels and may be safely used to investigate the role of Cdc42 in mouse sperm.

In addition to MLS-573151, other two specific pharmacological inhibitors were also used to study Cdc42. Secramine A blocks membrane recruitment of Cdc42 by impeding the release of GDI from Cdc42 (Cdc42/GDI complex), which sequesters the inactive GDP-bound Cdc42 in the cytosol and inhibits Cdc42 activation (GTP loading) (Pelish et al., 2006). CASIN targets Cdc42 by specifically blocking GEF activity on the Cdc42/GDI complex (referred to in (Peterson et al., 2006) as pirl1-related compound 2). On one hand, Secramine A partially impairs alkalinization-dependent CatSper activation, observed by a decrease in Em depolarization after NH_4_Cl addition in comparison with the control condition (without Cdc42 inhibitor) (Suppl. Fig. 5A-B). Moreover, the addition of Secramine A completely inhibited the alkalinization stimulated *I_CatSper_* (Suppl. Fig. 5C-D). On the other hand, because CASIN provoked a significant Em depolarization itself, it was not possible to observe the change in Em produced by NH_4_Cl (Suppl. Fig. 5E-F), however CASIN decreased the *I_CatSper_* partially (~50%) (Suppl. Fig. 5G-H) at negative and positive voltages. These findings show that both Cdc42 inhibitors have off-target effects on CatSper activity and are not suitable to evaluate the role of Cdc42 on mouse sperm physiology.

### Cdc42 is necessary for cAMP production by sAC

Our results identify Cdc42 as a new endogenous regulator of CatSper. Recently, it was demonstrated that CatSper activity is upregulated by a cAMP-dependent activation of PKA in mouse sperm (Orta et al., 2018). In the next set of experiments, we aimed to elucidate if Cdc42-dependent modulation of CatSper function occurs upstream PKA activation.

To investigate if Cdc42 inhibition alters the capacitation-induced PKA activation, PKA-dependent phosphorylation was evaluated in the presence or absence of Cdc42 inhibitor. Incubation with MLS-573151 led to a decrease in the phosphorylation of PKA substrates (pPKAs) and tyrosine residues (pY) (Fig. 7A). These data indicate that active Cdc42 is crucial for proper PKA activity. To analyze whether the requirement of Cdc42 is upstream PKA activation, sperm were incubated with Cdc42 inhibitor in the presence or absence of a membrane permeable analog of cAMP (8-Br-cAMP) and a phosphodiesterase inhibitor (IBMX). As a result, addition of 8-Br-cAMP/IBMX bypassed the inhibition of pPKAs and pY caused by 20 μM MLS-573151 (Fig. 7B) suggesting that Cdc42 activity is required upstream PKA activation. Furthermore, the decreased in [Ca^2+^]_i_ provoked by MLS-573151 was also bypassed by 8-Br-cAMP/IBMX (Fig. 7C and Suppl. Fig. 6A).

**Figure 7:**
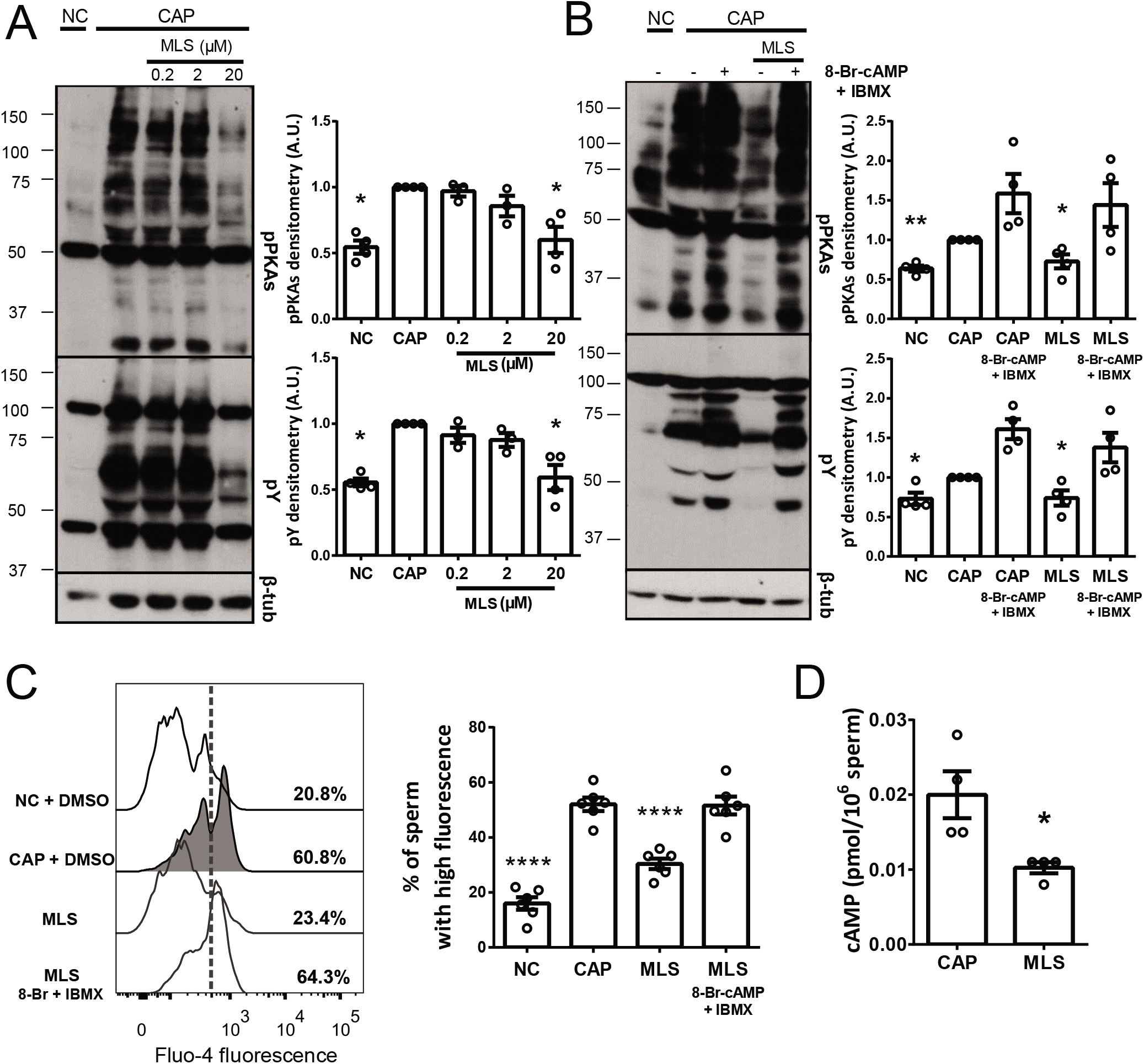
cAMP/PKA pathway inhibition was bypassed by using membrane permeable analogs of cAMP. **A-B)** Protein extracts were separated by 10% SDS-PAGE and immunoblotted with antibodies against phosphotyrosine residues (pY) and substrates phosphorylated by PKA (pPKAs). As a loading control, anti-β-tubulin was used. Sperm were incubated for 90 min under non-capacitating (NC) and capacitating conditions (CAP) in the absence (DMSO) or presence of the appropriate inhibitor. **A)** Increasing amounts of MLS-573151 produced a concentration-dependent decrease in pPKAs and pY phosphorylation. Quantitative analysis was performed by measuring the optical density of all bands and relativized to β-tubulin. Values represent the mean ± SEM of at least 3 independent experiments, where normalization to the control condition (CAP 90 min with DMSO) was used. * p<0.05 represents statistical significance vs. control (CAP 90 min with DMSO). Non-parametric Kruskal-Wallis test was performed in combination with Dunn’s multiple comparisons test. **B)** Sperm were incubated under capacitating conditions with 20 μM MLS-573151. The inhibition observed in this concentration was bypassed by 1 mM 8-Br-cAMP + 0.2 mM IBMX. Quantitative analysis was performed as described above. Values represent the mean ± SEM of 4 independent experiments, where normalization to the control condition (CAP 90 min with DMSO) was used. ** p<0.01, * p<0.05 represents statistical significance vs. CAP 90 min + 8-Br-cAMP/IBMX. Non-parametric Kruskal-Wallis test was performed in combination with Dunn’s multiple comparisons test. **C)** Ca^2+^ was measured in mouse sperm incubated for 90 min under non-capacitating (NC) or capacitating conditions (CAP) in the absence (DMSO) or presence of Cdc42 inhibitor. The inhibition observed with 20 μM MLS-573151 was bypassed by using 1 mM 8-Br-cAMP + 0.2 mM IBMX. Representative histograms of normalized frequency vs. Fluo-4 fluorescence of non-PI stained sperm (live), with the corresponding percentage of sperm that increased Fluo-4 fluorescence, are shown (left panel). The percentage of sperm that responds by increasing the [Ca^2+^]_i_, was established in the CAP control condition and extrapolated to the others conditions (dashed line). Percentage of sperm that increased Fluo-4 fluorescence after being incubated for 90 min in the different conditions (right panel). Values represent the mean ± SEM of 6 independent experiments. **** p<0.0001 represents statistical significance vs. control (CAP 90 min with DMSO). One-way ANOVA with Dunnett’s multiple comparisons test was performed. **D)** Total cAMP content of sperm incubated for 90 min under capacitating conditions with or without 20 μM MLS-573151. Values represent the mean ± SEM of 4 independent experiments. * p<0.05 represents statistical significance vs. control (CAP 90 min with DMSO). Unpaired t-test was performed.

Since our results indicate that Cdc42 is acting upstream PKA activity, we analyzed whether active Cdc42 is necessary for cAMP production by sAC. Sperm cAMP levels were measured in the presence or absence of 20 μM MLS-573151 and as a result, a significant decreased in cAMP concentration was observed after incubation with the Cdc42 inhibitor (Fig. 7D). All together, these results suggest that Cdc42 activity is required for cAMP production by sAC, and consequently for the upregulation of CatSper through PKA.

We also confirmed that the other Cdc42 inhibitors, that were discarded in previous experiments (Suppl. Fig. 5), display non-specific inhibition on CatSper function. Both Secramine A and CASIN provoke a decrease in phosphorylation patterns of PKAs and pY (Suppl. Fig. 6B) that could be restored by 8-Br-cAMP/IBMX (Suppl. Fig. 6C), but contrary to MLS-573151, sustained [Ca^2+^]_i_ inhibition was not recovered even with the addition of membrane permeable analog of cAMP and IBMX (Suppl. Fig. 6D-G). This reinforces that these drugs are not suited to study Cdc42 function in mouse sperm.

Finally, to further support our approach, the CatSper-dependent Ca^2+^ increase was investigated in the presence of PKA and CatSper inhibitors, where the addition of 8-Br-cAMP/IBMX should not be able to restore Ca^2+^ levels. In one case, because CatSper is stimulated downstream PKA activation and hence, direct inhibition of the kinase is not bypassed by cAMP analogs. In the other case, the direct blockade of the channel with CatSper inhibitors also impedes its cAMP-dependent activation. In agreement with our interpretation of the results presented above, the decrease in the [Ca^2+^]_i_ promoted by either PKA inhibition (KT-5720) or CatSper blockage (NNC55-0396 or Mibefradil) was not restored by PKA activation using 8-Br-cAMP/IBMX (Suppl. Fig. 7A-C).

## Discussion

Cdc42 is a member of the Rho-family of small GTPases which regulate several cellular functions such as gene transcription, actin cytoskeleton organization, cell motility, polarity, growth, and survival (Hall, 2012). Cdc42 was observed in the acrosomal region and flagella of several mammalian sperm (Baltiérrez-Hoyos et al., 2012; Delgado-Buenrostro et al., 2005; Ducummon and Berger, 2006). With the exception of one report that described the involvement of Cdc42 in acrosomal exocytosis (Baltiérrez-Hoyos et al., 2012), its role has not been conclusively demonstrated, in particular, given the fact that knocking out the *Cdc42* gene results in embryonic lethality (Chen et al., 2000). In this report, we identified a new function of the protein Cdc42 in mouse sperm. By localizing in the CatSper signaling complex, Cdc42 activity is necessary for cAMP production which in turn, alters the observed upregulation of CatSper by PKA.

By using super-resolution microscopy, the small GTPase Cdc42 was localized in the sperm flagellum forming four columns along the principal piece, resembling the localization of CatSper. Additionally, CatSper1 KO sperm displayed a disorganized Cdc42 distribution in the principal piece. Previous reports demonstrate that CatSper scaffolds P-CaMKII, Caveolin-1 and PP2B in this defined quadrilateral structure, as sperm from CatSper1-null mice showed these proteins delocalized becoming uniformly distributed in the plasma membrane (Chung et al., 2014). Therefore, our results support the idea that Cdc42 might be a new component of these highly organized Ca^2+^ domains. Until now, there is no direct evidence of these molecules regulating CatSper activity. On the contrary, Caveolin-1-null mice are fertile (Razani et al., 2001) and CatSper channels in these sperm are localized normally (Chung et al., 2014). Furthermore, PP2B-null mice are infertile, as their sperm cannot develop hyperactivation, which results in deficiencies to penetrate the zona pellucida. Although this evidence might suggest CatSper impairment, no defects in the principal piece were observed in PP2B-null mice, but instead these sperm exhibit a rigid mid piece (Miyata et al., 2015).

Previous reports described the interaction between Cdc42 and Caveolin-1. Both proteins were co-immunoprecipitated from guinea pig and mouse sperm (Baltiérrez-Hoyos et al., 2012). The similar domain organization observed for both Cdc42 and Caveolin-1 may indicate possible functional interactions between these two proteins, although this needs to be further investigated.

Here we propose Cdc42 as a new endogenous modulator of CatSper activity in the Ca^2+^ signaling domains orchestrated by this channel. This discovery has important implication for understanding the molecular regulation of CatSper. In this regard, recent reports found that mouse CatSper, is upregulated by a cAMP-dependent activation of PKA (Orta et al., 2018). However, it is still unknown whether PKA phosphorylates one or more subunits of the channel or if its activation relays on other intermediary events. In this report, we obtain similar results in terms of CatSper function and PKA activity. However, we are proposing a new regulatory mechanism that connects CatSper function and the cAMP/PKA pathway. We found that Cdc42 is important for cAMP production by sAC, which in turn results in activation of PKA. We hypothesize that the localization of Cdc42 in the CatSper signaling complex is essential for the tight interplay between the increase in the [Ca^2+^]_i_, and the cAMP/PKA pathway. Although a cross talk between cAMP/PKA-dependent pathways and Ca^2+^ clearly plays a key role in sperm capacitation, the connection between these signaling events is incompletely understood and our results contribute to unravel this complex regulatory mechanism.

Intracellular alkalinization has been univocally demonstrated to alter CatSper function endogenously (Kirichok et al., 2006). In this regard, it was recently reported the asymmetrically distribution of Hv1 channel in the human sperm flagellum, which is responsible, at least in part, for the increase in pH_i_, (Lishko et al., 2010). This specific localization of Hv1 could produce local alkalization when positioned in close proximity to a subset of CatSper channels resulting in asymmetric local Ca^2+^ changes that may be required for the complex movement of the sperm flagellum (Miller et al., 2018). In mouse, the alkalinization of the cytoplasm during capacitation occurs as a result of Na^+^-H^+^ exchangers (NHE) (Chávez et al., 2014; Wang et al., 2003; Wang et al., 2007) since there is no clear evidence showing a significant role of Hv1 channels. Recent reports demonstrated that EFCAB9 is a pH and Ca^2+^ dependent sensor that drives the activation of CatSper (Hwang et al., 2019). It is also possible that the highly histidine rich amino terminus of CatSper1 protein is also involved in sensing pH_i_, (Ren et al., 2001) since EFCAB9-null sperm display a modest CatSper activation in response to higher pH_i_, (Hwang et al., 2019). Modulation of CatSper by pH_i_, and Cdc42 might be linked since a connection between Cdc42 and H^+^ efflux through NHE1 has been reported in somatic cells. A bistable positive feedback regulation between Cdc42 and NHE1 activities has been previously proposed (Frantz et al., 2007).

Rho GTPases act by a tightly regulated switch between their inactive GDP-bound and their active GTP-bound state. The later elicits its actions through target proteins. To date, a testis-specific Rho target protein Rhophilin has been described, which interacts with Ropporin (Fujita et al., 2000). Ropporin is mainly localized in sperm flagellum and binds A-kinase anchoring protein 3 (AKAP3) (Carr et al., 2001) through cAMP-dependent mechanism. Some reports suggest a role of Rho/Rhophilin/Ropporin in sperm motility regulation, as it was demonstrated that Ropporin expression is diminished in sperm from asthenozoospermic samples (Chen et al., 2011).

In summary, in this study we present novel findings indicating that Cdc42 plays a central role in mammalian sperm capacitation. Figure 8 summarizes the results of this work in a proposed model. In the female reproductive tract (or when sperm are exposed to conditions that support capacitation), HCO_3_^-^stimulation of sAC resulting in cAMP production requires the activity of Cdc42. This protein is localized in the signaling complex organized by CatSper together with other proteins (not shown in this simplified model). The increase in cAMP levels promotes the activation of PKA which in turn, phosphorylates a subset of proteins that lead to phosphorylation in tyrosine residues of others. In addition, the stimulation of the cAMP/PKA pathway leads to the activation of CatSper (either directly or through other intermediates). As a result, a sustained Ca^2+^ influx promotes the development of hyperactivation. Considering that CatSper is sperm specific and has a fundamental role in hyperactivation, this channel has emerged as a potential therapeutic target in male infertility as well as in the development of contraceptive strategies. Thus, understanding the modulation of CatSper activity and all the players involved in its regulation is of great importance.

**Figure 8:**
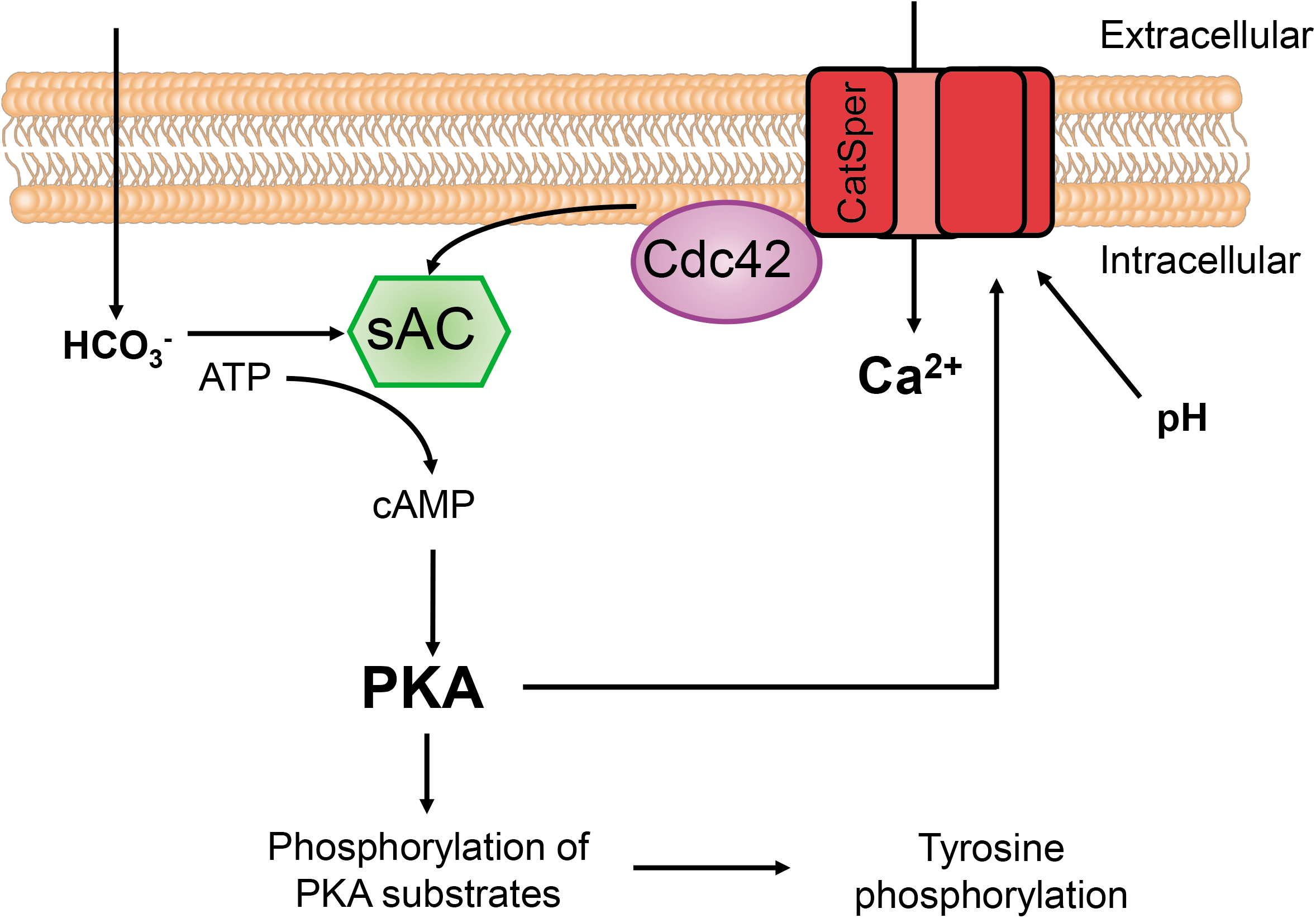
Proposed model for CatSper activation during mouse sperm capacitation. Cdc42 is localized in the signaling complex organized by CatSper together with other proteins (not shown in this simplified model). Cdc42 is necessary for the HCO_3_^-^stimulation of sAC, resulting in cAMP production and activation of PKA. This activation provokes the phosphorylation of substrates (by PKA or in tyrosine residues) and the activation of CatSper (either directly or through other intermediates). As a result, a sustained Ca^2+^ influx promotes the development of hyperactivation.

## Materials and Methods

### Reagents

Chemicals were obtained from the following sources: bovine serum albumin (BSA) A7906, 3-isobutyl-1-methylxanthine (IBMX), dibutyryl-cyclic AMP (db-cAMP), Mibefradil, NNC55-0396, anti-β-tubulin T4026, protease inhibitor cocktail P8340, carbonyl cyanide m-chlorophenylhydrazone (CCCP), guanosine 5’-triphosphate sodium salt hydrate (GTP) G8877 and ethylene glycol-bis (2-aminoethylether)-N,N,N,N’tetraacetic acid (EGTA) were purchased from Sigma-Aldrich (St. Louis, MO, USA). Anti-phosphotyrosine (anti-pY) clone 4G10 and anti-facilitative glucose transporter 3 (anti-GLUT3) AB1344 from Millipore (Temecula, CA, USA); anti-phospho PKA substrates (anti-pPKAs) clone 100G7E and anti-phospho CaMKII (Thr286) 3361 from Cell Signaling Technology (Danvers, MA, USA); horseradish peroxidase-conjugated (HRP) anti-rabbit IgG and HRP anti-mouse IgG from Vector Laboratories (Burlingame, CA, USA). 2’,7’-Bis-(2-Carboxyethyl)-5-(and-6)-Carboxyfluorescein, acetoxymethyl ester (BCECF AM), 8-Bromo-cyclic AMP (8-Br-cAMP, sodium salt), KT-5720, CASIN and MLS-573151 from Cayman Chemicals (Ann Arbor, MI, USA). Secramine A was kindly provided from the Kirchhausen Lab (Harvard Medical School, MA, USA). Fluo-4 AM, 3,3-dipropylthiadicarbocyanine iodide (DiSC_3_(5)), pluronic acid, F(ab’)2-goat anti-rabbit IgG-Alexa Fluor 647 (A-21246), and chicken anti-mouse IgG-Alexa Fluor 647 (A-21463) from Invitrogen, Thermo Fisher Scientific (Waltham, MA, USA); while propidium iodide (PI), anti-Cdc42 (B-8) sc-8401, Cdc42 (B-8) Blocking Peptide sc-8401 P, anti-Cdc42 sc-87 and anti-Actin sc-1616-R from Santa Cruz Biotechnology (Dallas, TX, USA). Anti-AKAP82 611564 from BD Biosciences (Franklin Lakes, NJ, USA). Anti-β-tubulin E7 was obtained from Developmental Studies Hybridoma Bank (DBHS) University of Iowa; E7 was deposited to the DSHB by Michael Klymkowsky. All other chemicals were of reagent grade. Fluo-4 AM, DiSC_3_(5), pluronic acid, IBMX, KT-5720, CCCP, CASIN, MLS-573151 and Secramine A were dissolved in DMSO; EGTA, 8-Br-cAMP, db-cAMP, Mibefradil, NNC55-0396 and PI were dissolved in hexa-distilled water.

### Animals

Hybrid F1 (C57BL/6 male x Balb/C female) and CD-1 mature (10–12 weeks-old) male mice were used. CatSper1 KO (Ren et al., 2001) mice and their corresponding heterozygous (HET) siblings (C57BL/6) reside at the University of Massachusetts (Navarrete et al., 2016). CatSper1-null mice were euthanized in accordance with the Animal Care and Use Committee (IACUC) guidelines of UMass-Amherst (protocol #2016-0026). In all cases, mice were housed in groups of 4 or 5 in a temperature-controlled room (23°C) with lights on at 07:00 h and off at 19:00 h, and had free access to tap water and laboratory chow. All experimental procedures were carried according to guidelines of the institutional animal care and were reviewed and approved by the Ethical Committees of the *Instituto de Biología y Medicina Experimental, Buenos Aires* and of the *Instituto de Biotecnología, Cuernavaca.* Experiments were performed in strict accordance with the Guide for Care and Use of Laboratory Animals approved by the National Institutes of Health (NIH).

### Sperm medium

The non-capacitating medium used in this study was a modified Toyoda–Yokoyama–Hosi (modified TYH) containing 119.3 mM NaCl, 4.7 mM KCl, 1.71 mM CaCl_2_.2H_2_O, 1.2 mM KH_2_PO_4_, 1.2 mM MgSO_4_.7H_2_O, 0.51 mM sodium pyruvate, 5.56 mM glucose, 20 mM HEPES and 10 μg/ml gentamicin (NC medium). For capacitating conditions 15 mM NaHCO_3_ and 5 mg/ml BSA were added (CAP medium). In all cases, pH was adjusted to 7.4 with NaOH. In *in vitro* fertilization assays, 20 mM HEPES was omitted while 25 mM NaHCO_3_ and 4 mg/ml BSA were added (TYH IVF).

### Sperm capacitation

Animals were euthanized and cauda epididymal mouse sperm were collected. Both cauda epididymis were placed in 1 ml of non-capacitating modified TYH medium (NC: without BSA and NaHCO_3_) in the presence or absence of Ca^2+^ as described for each experiment. After 15 min of incubation at 37°C (swim-out), epididymis were removed, and sperm were resuspended to a final maximum concentration of 1×10^7^ cells/ml on 100 μl of the appropriate medium, depending on the experiment performed.

A 10 min pre-incubation in 100 μl of NC medium containing Cdc42 inhibitors (MLS-573151, CASIN or Secramine A), CatSper inhibitors (30 μM Mibefradil or 10 μM NNC55-0396) or PKA inhibitor (30 μM KT-5720) was conducted when required. An equal volume (100 μl) of NC or two-fold concentrated capacitating medium (CAP 2×: 30 mM NaHCO_3_ and 10 mg/ml BSA) containing the appropriate inhibitors was added. When necessary, 1 mM 8-Br-cAMP or db-cAMP (permeable cAMP analogs) in combination with 0.2 mM IBMX (phosphodiesterase inhibitor) were also added. Finally, sperm were incubated for 90 min at 37°C.

### Three-Dimensional Stochastic Optical Reconstruction Microscopy (3D STORM)

After swim-out, cells were washed twice with NC medium by centrifugation (5 min at 400 x *g*) and finally resuspended in NC medium. Sperm were seeded in poly-L-lysine coated coverslips (Corning #1.5), air-dried for 10 min, fixed with 4% fresh paraformaldehyde in PBS for 10 min at room temperature, and followed by three washes with PBS (5 min at 400 x *g*). Cells were then permeabilized with 0.5% Triton X-100 in PBS for 5 min at room temperature and washed three times with PBS (5 min at 400 x *g*). Samples were blocked with 3% BSA/PBS for 1 h at room temperature and then incubated overnight at 4°C with primary antibody in a humidified chamber: anti-Cdc42 (1:100), anti-β-tubulin E7 (1:100), anti-AKAP4 (1:300), anti-GLUT3 (1:100) or anti-P-CaMKII (1:50), all diluted in 1% BSA/PBS. Cells were washed three times with T-PBS (5 min at 400 x *g*), and further incubated with Alexa Fluor 647-conjugated anti-mouse (1:500) or rabbit (1:1000) secondary antibody diluted in 1% BSA/PBS for 1 h at room temperature. Sperm were then washed with T-PBS twice and PBS once for 5 min each, and immediately mounted in STORM imaging buffer (50 mM Tris-HCl pH 8, 10 mM NaCl, 0.56 mg/ml glucose oxidase, 34 μg/ml catalase, 10% glucose, and 1% β-mercaptoethanol). Nonspecific staining was determined by incubating the sperm in the absence of primary antibody. Images were acquired using Andor IQ 2.3 software in a custom built microscope equipped with an Olympus PlanApo 100x NA 1.45 objective and a CRISP ASI autofocus system (Campagnola et al., 2015; Weigel et al., 2011). Alexa Fluor 647 was excited with a 642 nm laser (DL640-150-O, CrystaLaser, Reno, NV) under continuous illumination. Initially the photo-switching rate was sufficient to provide a substantial fluorophore density. However, as fluorophores irreversibly photo-bleached, a 405 nm laser was introduced to enhance photo-switching. The intensity of the 405 nm laser was adjusted in the range 0.01–0.5 mW to maintain an appropriate density of active fluorophores. Axial localization was achieved via astigmatism using a MicAO 3DSR adaptive optics system (Imagine Optic, Orsay, France) that allows both the correction of spherical aberrations and the introduction of astigmatism (Izeddin et al., 2012). Axial localization with adaptive optics enabled 3D reconstruction over a thickness of 1 μm (Gervasi et al., 2018). A calibration curve for axial localization was generated with 100 nm TetraSpeck microspheres (Invitrogen) immobilized on a coverslip (Huang et al., 2008). The images were acquired in a water-cooled, back-illuminated EMCCD camera (Andor iXon DU-888) operated at −85°C at a rate of 23 frames/sec. 50,000 frames were collected to generate each super-resolution image. Single-molecule localization, drift correction using image cross-correlation and reconstruction were performed with Thunder STORM (Ovesný et al., 2014).

To find the molecular radial distributions, we selected regions of interest of the flagellum that were found to lie in a straight line. The center of the flagellar cross-section was first found by Gaussian fitting of the localization histograms along x and y. The coordinates of the localized molecules were then transformed into cylindrical coordinates to obtain the radial position *r* and azimuthal angle θ (Gervasi et al., 2018; Stival et al., 2018).

### Determination of [Ca^2+^]_i_ and pH_i_ by flow cytometry

Sperm [Ca^2+^]_i_, and pH_i_, changes were assessed using Fluo-4 AM and BCECF AM respectively as previously described (Luque et al., 2018). After incubation in the appropriate medium, samples were centrifuged at 400 x *g* for 4 min at room temperature and resuspended in 200 μl of NC medium containing either 1 μM Fluo-4 AM and 0.02% pluronic acid or 0.5 μM BCECF AM for 20 min at 37°C. Samples were washed again and resuspended in 50 μl of NC medium. Before collecting data 2 ng/μl of PI was added to monitor viability. Data were recorded as individual cellular events using a BD FACSCanto II TM cytometer (Biosciences; Becton, Dickinson and Company). Side-scatter area (SSC-A) and forward-scatter area (FSC-A) data were collected from 20000 events per sample in order to define sperm population as previously described (Escoffier et al., 2012). In all cases, doublet exclusion was performed analyzing two-dimensional dot plot FSC-A vs. forward-scatter height (FSC-H). Positive cells for Fluo-4 AM were collected using the filter for Fluorescein isothiocyanate (FITC; 530/30), and for PI the filter for Peridinin chlorophyll protein complex (PerCP; 670LP). The two indicators had minimal emission overlap, but still compensation was done. Data were analyzed using FlowJo software (V10.0.7).

### Cdc42 activation assay

To specifically detect the GTP-loaded form of Cdc42, the Cd42 G-LISA kit (Cytoskeleton Inc., Denver, CO, USA) was used according to the manufacturer’s instructions. Briefly, 50 μl of duplicate samples of lysates from sperm incubated for 90 min under capacitating conditions (CAP) in the absence (DMSO) or presence of Cdc42 inhibitor (20 μM MLS-573151) at a concentration between 0.65 and 0.95 mg protein/ml were added into each well. Plain lysis buffer and a standard of constitutively active purified GTP-bound Cdc42 protein were added to duplicate wells as a blank and a positive control respectively. After binding, anti-Cdc42 primary antibody was added to each well followed by secondary antibody labeled with HRP, which was developed by adding HRP reagent. Each well was read at OD 492 nm on a 96-well plate spectrophotometer. Lysis buffer background was subtracted and results were normalized to protein concentration.

### Membrane potential assay in cell populations

The procedure was done as previously described (Demarco et al., 2003). Sperm obtained from swim-out were diluted in NC or CAP medium and loaded with 1 μM of the membrane-potential-sensitive dye DiSC_3_(5) for 5 min. Mitochondrial membrane potential was dissipated incubating sperm for 2 min with 500 nM CCCP. After this, 1 ml of the sperm suspension was transferred to a stirred cuvette at 37°C and fluorescence monitored with an Ocean Optics USB4000 spectrofluorometer operated by Spectra Suite (Ocean Optics, Inc., USA) at 620/670 nm excitation/emission wavelength pair (López-González et al., 2014). Cell hyperpolarization decreases the dye fluorescence. Recordings were initiated after reaching steady-state fluorescence (1-3 min) and calibration was performed at the end of each measure by adding 1 μM valinomycin and sequential additions of KCl as previously described (Demarco et al., 2003). Sperm membrane potential (Em) was obtained from the initial fluorescence (measured as Arbitrary Fluorescence Units) by linearly interpolating it in the theoretical Em values for the calibration curve against arbitrary fluorescence units of each trace. This internal calibration for each determination compensates for variables that influence the absolute fluorescence values.

On one hand, CatSper activity was assessed measuring Em in TYH medium after adding EGTA (3.5 mM final, pH adjusted with NaOH to ~10 so that media pH does not change upon H^+^ release in exchange for Ca^2+^), which allows Na^+^ influx through CatSper, depolarizing non-capacitated sperm. The magnitude of the depolarization caused by Na^+^ influx correlates with the extent of CatSper opening. Briefly, after sperm treatment (capacitation in presence or absence of Cdc42 inhibitors), cells were collected by centrifugation (400 x *g*, 3 min) and concentration adjusted to 2×10^6^ sperm/ml with NC medium with the addition of the corresponding inhibitor or not. Then, sperm were loaded with 1 μM of DiSC_3_(5). Results were represented as ΔFluorescence, the difference between DiSC_3_(5) fluorescence after 3.5 mM EGTA addition and before (resting Em) (Ernesto et al., 2015). In these experiments, calibration curves were performed to confirm that sperm were viable. Em values were not calculated from these calibrations.

On the other hand, non-capacitated sperm were loaded with 1 μM of DiSC_3_(5) and then exposed to 10 mM NH_4_Cl, with prior addition of Cdc42 inhibitor or not. Alkalinization allows CatSper activation, which results in Ca^2+^ influx depolarizing the cell. Again, the magnitude of the depolarization correlates with the extent of CatSper opening. Results were represented as ΔEm, the difference between Em after 10 mM NH_4_Cl addition and before (Em with or without Cdc42 inhibitor).

### Electrophysiology

Cauda epididymal sperm obtained from swim-out were resuspended in HS medium containing: NaCl 135 mM, KCl 5 mM, CaCl_2_ 2 mM, MgSO_4_ 1 mM, lactic acid 10 mM, piruvic acid 1 mM, glucose 5 mM, HEPES 20 mM, pH 7.4 (NaOH). One hundred μl aliquots of the cell suspension were dispensed into a recording chamber (1 ml total volume) and subjected to electrophysiological recording. Whole-cell macroscopic CatSper monovalent currents (*I_CatSper_*) were obtained by patch-clamping the sperm cytoplasmic droplet in cauda epididymal sperm and were analyzed as reported previously (Ernesto et al., 2015; Kirichok et al., 2006). Seals between the patch pipette and the cytoplasmic droplet in sperm were formed in HS bath solution (baseline current). After establishing the whole-cell configuration, the bath solution was changed for divalent cation free solution (DVF) which is best to measure CatSper-dependent currents. The DVF bath solution contained: Na-gluconate 150 mM, Na_2_EDTA 2 mM, EGTA 2 mM, HEPES 20 mM, pH 7.4, while the pipette solution contained: Met-Cs 135 mM, CsCl 5 mM, Na-ATP 5 mM, EGTA 10 mM, HEPES 10 mM, pH 7.0. When required, 0.4 mM of GTP was added to the patch-clamp pipette solution. The osmolarity of all solutions was adjusted with dextrose. All recordings were performed using patch-clamp amplifier (Axopatch 200 Molecular Devices) at room temperature (22°C). Pulse protocols and data capture were performed using pCLAMP6 software (Molecular Devices); data analysis was carried out with Clampfit 10.6 (Molecular Devices), Origin 7.5 (Microcal Software) and Sigma Plot 10 (SYSTAT Software). Current records, unless indicated otherwise, were acquired at 20–100 kHz and filtered at 5–10 kHz (low-pass Bessel filter) using a computer attached to a DigiData 1200 (Molecular Devices). Patch pipettes were pulled from borosilicate glass (Kimble® Queretaro) and had a final resistance between 15–20 MΩ. *I_CatSper_* currents were evoked employing a conventional voltage-ramp protocol from −80 mV to +80 mV, with duration of 750 ms from a holding potential of 0 mV. In all cases, the addition of Cdc42 inhibitors was recorded until reaching a stable-state effect (3-5 min).

### Extraction of sperm proteins and western blotting

After incubation in the appropriate medium, sperm were washed by centrifugation (5 min at 400 x *g*), resuspended in sample buffer without reducing agents (62.5 mM Tris–HCl pH 6.8, 2% SDS, 10% glycerol) and boiled for 5 min. After centrifugation for 5 min at 13,000 x *g*, 5% β-mercaptoethanol and 0.0005% bromophenol blue was added to the supernatants then boiled again for 5 min. Protein extracts equivalent to 2-4×10^6^ sperm per lane were separated by SDS-PAGE in gels containing 10% polyacrylamide and transferred onto nitrocellulose membranes. For Cdc42 detection, after being washed sperm were incubated in RIPA buffer (50 mM Tris, 150 mM NaCl, 0.5% Sodium deoxycholate, 1% NP-40, 0.1% SDS) with protease inhibitor cocktail 1X for 20 min on ice, and proteins from 10-14×10^6^ sperm per lane were separated by SDS-PAGE in 15% polyacrylamide gels. In all cases, blots were blocked in 5% nonfat dry milk in PBS containing 0.1% Tween 20 (T-PBS) for 1 h at room temperature. An overnight incubation at 4°C with the primary antibody anti-Cdc42 was required, while 1 h incubation at room temperature with the other primary antibodies was sufficient. Antibodies were diluted in 2% nonfat dry milk in T-PBS as follows: 1:500 for anti-Cdc42 (Santa Cruz sc-8401); 1:750 for anti-Cdc42 (Santa Cruz sc-87); 1:3000 for anti-pPKAs, anti-pY and anti-β-tubulin T4026. The corresponding secondary antibodies were incubated for 1 h at room temperature, diluted in 2% nonfat dry milk in T-PBS as follows: 1:3000 for HRP anti-rabbit and 1:3000 for HRP anti-mouse. In all cases the reactive bands were visualized using a chemiluminescence detection solution consisting of 100 mM Tris-HCl buffer pH 8, 205 μM coumaric acid, 1.3 mM luminol, 0.01% H_2_O_2_ and were exposed for different time periods to CL-XPosure film (Thermo Scientific). In all experiments, molecular masses were expressed in kiloDaltons (kDa). ImageJ 1.48k (National Institute of Health, USA) was used for analysis of the western blot images following the specifications of ImageJ User Guide, IJ 1.46r. The optical density of all bands were quantified and normalized, first to the β-tubulin or Actin band and then to the capacitating condition of each experiment.

### Immunofluorescence

To perform immunofluorescence on mouse sperm, a previously described method was used (Gervasi et al., 2018). Briefly, mouse sperm were washed twice by centrifugation for 5 min at 400 x *g*, resuspended in PBS containing 4% paraformaldehyde (Electron Microscopy Sciences, Hatfield, PA, USA) for 10 min at room temperature. After washing twice for 5 min in PBS sperm were placed onto glass slides. Sperm were air-dried and then permeabilized with 0.5% Triton X-100 in PBS for 5 min. The slides were washed twice for 5 min in T-PBS and blocked in 3% BSA in PBS for 1 h at room temperature, then were incubated with primary antibody (1:100 mouse monoclonal anti-Cdc42 (B-8), sc-8401) diluted in PBS containing 1% BSA overnight at 4°C. Following a washing step (three times, 5 min in T-PBS), slides were incubated with Alexa Fluor 488-conjugated goat anti-mouse IgG (Invitrogen, A11001) diluted 1:500 in PBS containing 1% BSA for 1 h at room temperature, washed again for three times in T-PBS, and mounted using Vectashield mounting media (H-1000, Vector Labs, Burlingame, CA, USA). Nonspecific staining was determined by incubating the sperm with blocked antibody. To this end, the primary antibody (1:100 anti-Cdc42 (B-8), sc-8401) was pre-incubated with 20x of blocking peptide (Cdc42 (B-8), sc-8401 P) in PBS, for 2 h at room temperature with agitation. Slides were examined using an inverted fluorescence microscope (Nikon TE300) and images were captured at 60x magnification (Plan Apo, NA1.4(oil)), with a sCMOS camera (Andor, Zyla).

### Determination of cAMP levels

Intracellular sperm cAMP concentrations were determined using a cAMP select ELISA kit (Cayman Chemicals, Ann Arbor, MI, USA). After incubation of sperm in the appropriate conditions, reactions were stopped with 0.1 M HCl and cells were lysed with Triton 1%. This ELISA based assay has an increased sensitivity that ranges from 0.02 pmol to 40 pmol of cAMP. A standard curve was run for each assay, and the unknown cAMP concentrations were obtained by interpolation as recommended by the manufacturer.

### Sperm motility analysis

After incubation in the appropriate medium, sperm suspensions were loaded on a 100 μm chamber slide (Leja Slide, Spectrum Technologies) and placed on a microscope (Nikon Eclipse E200) stage at 37°C coupled to a BASLER acA780-75gc camera. Sperm movements were examined using computer-assisted semen analysis (CASA) system (Sperm Class Analyzer: SCA evolution, Microptic). Parameters used were as follows: 30 frames acquired, frame rate of 60 Hz, and cell size of 30-170 μm^2^. At least 20 microscopy fields corresponding to a minimum of 200 sperm were analyzed in each experiment. The following parameters were measured: mean path velocity (VAP), curvilinear velocity (VCL), straight line velocity (VSL), linearity (LIN), amplitude of lateral head displacement (ALH), and straightness (STR). Sperm were considered hyperactivated when presenting VCL ≥ 271 μm/sec, LIN < 50 %, and ALH ≥ 3.5 μm.

### In vitro fertilization assay

Eight to 12-week-old F1 female mice were superovulated using equine chorionic gonadotropin (5 IU, PMSG; Syntex) at 19:30, followed 48 h later by human chorionic gonadotropin (5 IU, hCG; OVUSYN, Syntex) intraperitoneal injection. Cumulus oocyte complexes (COC) were collected from oviducts 12–13 h after hCG administration, placed in TYH IVF medium (containing 25 mM NaHCO_3_ and 4 mg/ml BSA, with no HEPES addition), and inseminated *in vitro* with capacitated sperm at a final concentration of 1×10^6^ sperm/ml (capacitation in presence or absence of Cdc42 inhibitors). After coincubation for 4 h at 37°C with 5% CO2, eggs were washed three times and placed on drops containing TYH IVF at 37°C with 5% CO2. Fertilization was assessed by visualization of 2-cell embryos 20 h later.

### Statistical analysis

Data are expressed as mean ± standard error of the mean (SEM) of at least 3 independent experiments from different mice for all determinations. Statistical analyses were performed using the GraphPad Prism 6 software (La Jolla, CA USA). The differences between means of only 2 groups were analyzed using a t-test (motility parameters, △Em and 2-cell embryos). The non-parametric Kruskal-Wallis test was performed in combination with Dunn’s multiple comparisons test to analyze normalized median fluorescence intensity (MFI) of Fluo-4 and normalized *I_CatSper_* currents. One-way analysis of variance (ANOVA) with Dunnett’s multiple comparisons test was performed to analyze percentage of sperm with high Fluo-4 fluorescence. A probability (p) value of p<0.05 was considered statistically significant. Parametric or non-parametric comparisons were used as dictated by data distribution.

## Acknowledgments

We would like to thank Drs. Gabriel Rabinovich and Mariana Salatino for their assistance in the flow cytometry experiments. We also thank Rene Baron, Fortabat and Williams foundations. We are grateful to Dr. Kirchhausen (Harvard Medical School, MA, USA) for providing Secramine A and Clara Marin-Briggiler, Paula Balestrini, Martina Jabloñski and Liza Schiavi-Ehrenhaus for critically reading this manuscript.

## Author contributions

GML, XX, AR, MGG, GO, JLDVB, CS, TD performed experiments; GML, XX, MGG, GO, Diego K, NG analyzed data. GML, MGB prepared the manuscript with contributions of all other authors. AD, PEV, Dario K discussed results and contributed with ideas. GML and MGB designed the study.

## Supplementary Figure Captions

**Suppl. Figure 1:**
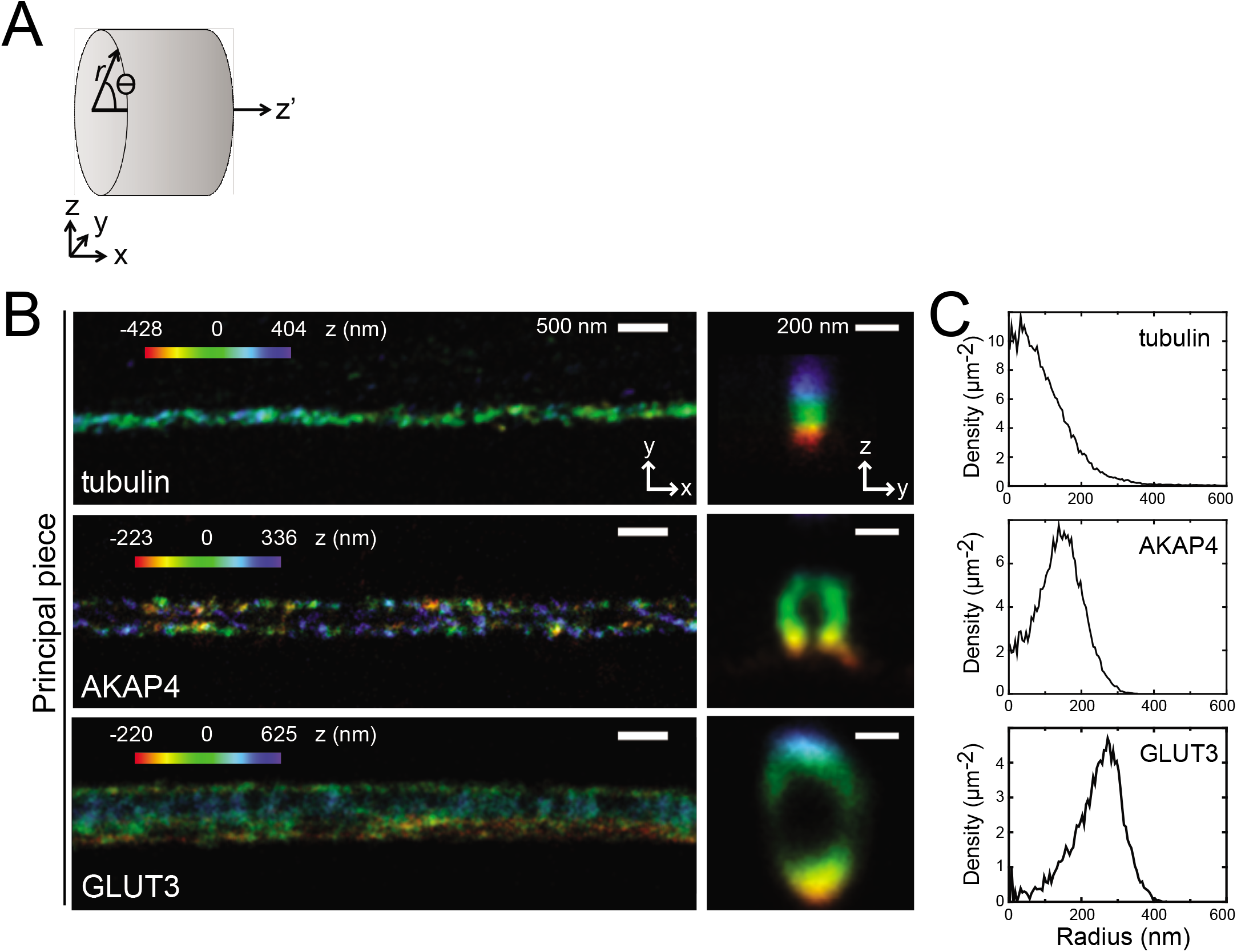
The structural information obtained by 3D STORM reproduces known protein localization in the sperm flagella. **A)** Cartoon of the cylindrical coordinate system for defining the radius and angles of molecular coordinates in STORM images. The longitudinal axis (x) is placed at the flagellar center and the origin at the annulus. **B)** Sperm were incubated under non-capacitating (NC) conditions. 3D STORM images of principal piece β-tubulin, AKAP4, and GLUT3. Left, x-y projections. Right, y-z cross-sections. **C)** Radial profiles of immunostained density of β-tubulin, AKAP4, and GLUT3. The color in all x-y projections encodes the relative distance from the focal plane along the z axis.

**Suppl. Figure 2:**
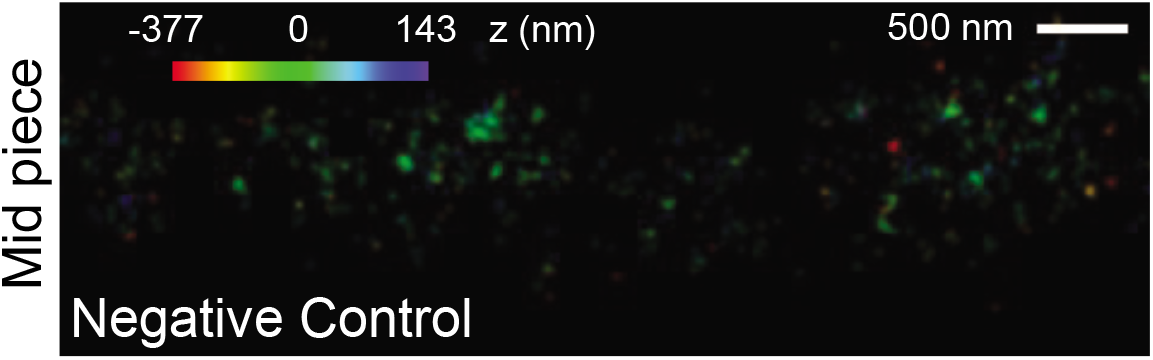
Sperm were incubated under non-capacitating (NC) conditions and negative controls were performed in the absence of primary antibody. 3D STORM images of mid piece, x-y projections. The color encodes the relative distance from the focal plane along the z axis.

**Suppl. Figure 3:**
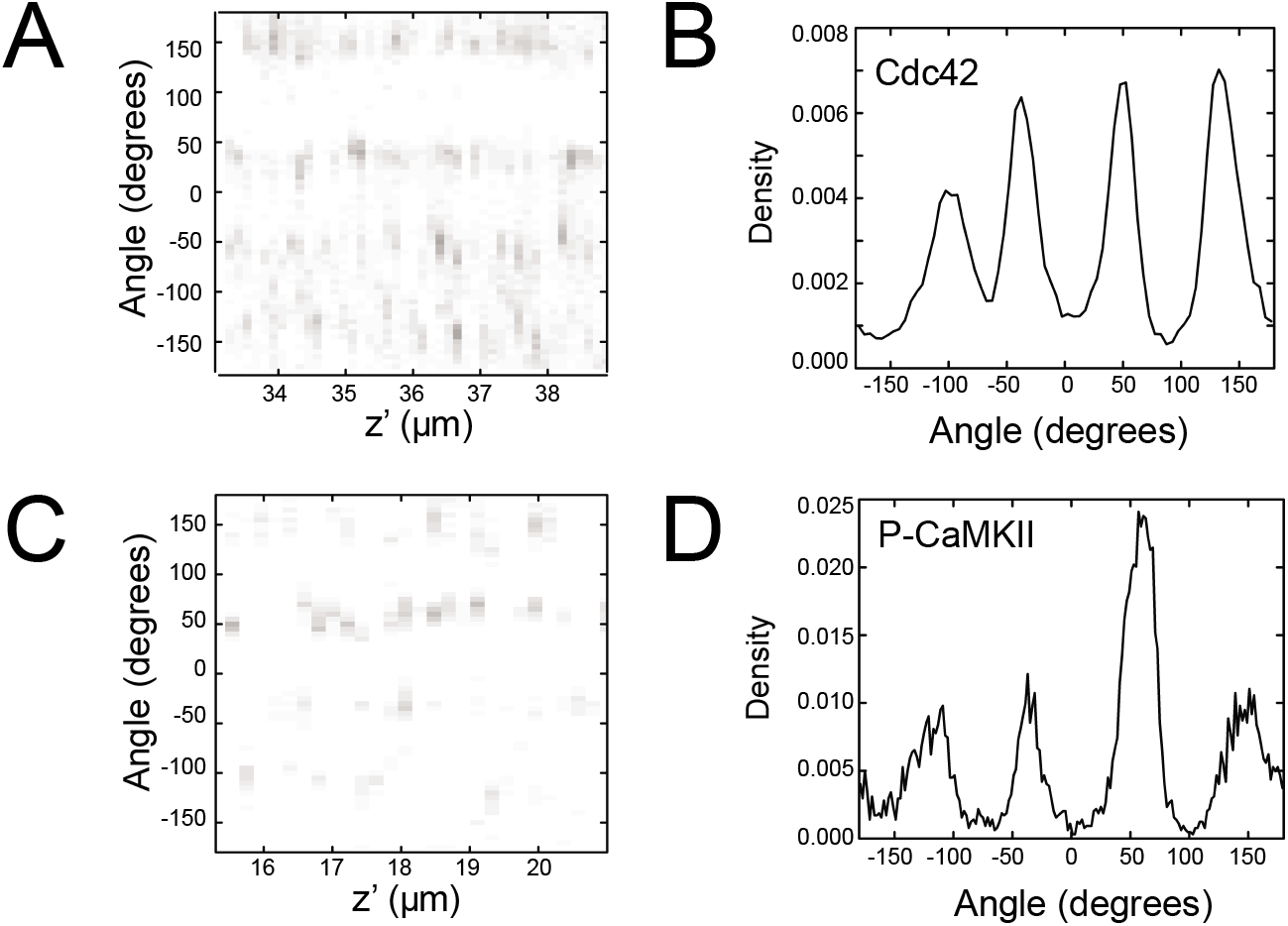
Cdc42 is localized in four longitudinal lines similar to P-CaMKII. **A)** Angular distributions of Cdc42. **B)** Profiles of the surface-localized Cdc42. **C)** Angular distributions of P-CaMKII. **D)** Profiles of the surface-localized P-CaMKII.

**Suppl. Figure 4:**
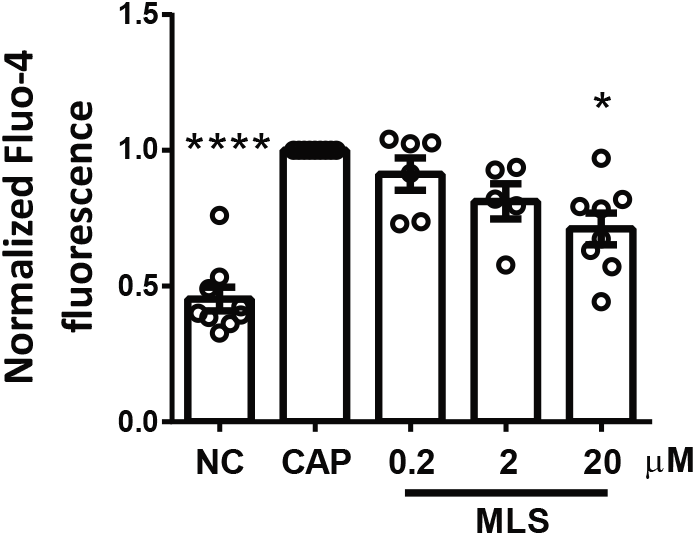
Cdc42 inhibition blocked the capacitation-associate increase in [Ca^2+^]_i_ in a dose dependent manner. Sperm were incubated for 90 min under non-capacitating (NC) and capacitating conditions (CAP) in the absence (DMSO) or presence of Cdc42 inhibitors: MLS-573151 (MLS), Secramine A and CASIN. Increasing μM concentrations of Cdc42 inhibitors were used, which resulted in a decrease in [Ca^2+^]_i_, in a dose dependent manner. Normalized MFI of Fluo-4 compared to the control condition (CAP 90 min with DMSO). Values represent the mean ± SEM of at least 6 independent experiments. **** p<0.0001, *** p<0.001 represents statistical significance vs. control (CAP 90 min with DMSO). Non-parametric Kruskal-Wallis test was performed in combination with Dunn’s multiple comparisons test.

**Suppl. Figure 5:**
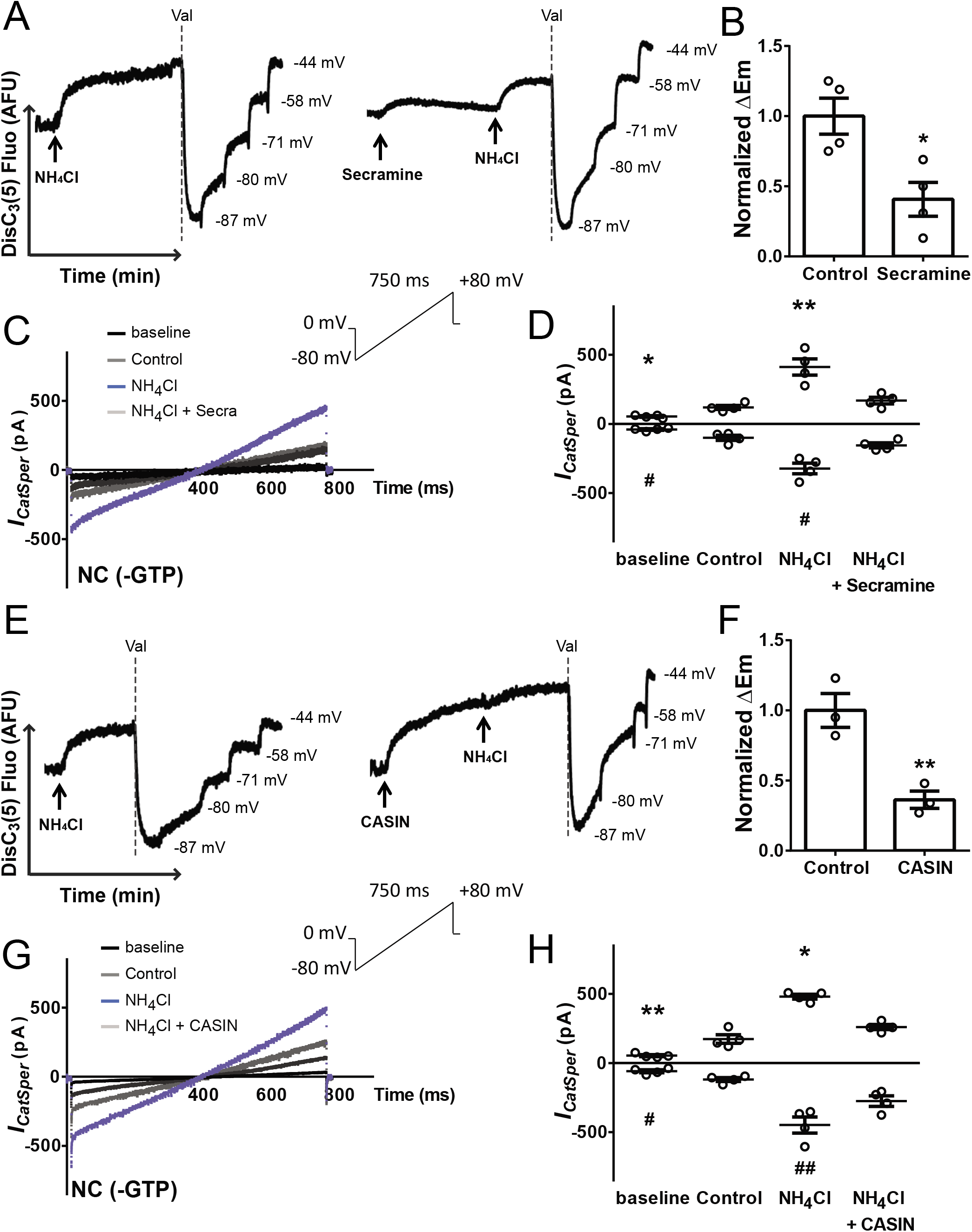
Alkalinization-induced depolarization responds differently to pre-incubation with Cdc42 inhibitors. CatSper activity was analyzed by measuring the Em of sperm obtained from swim-out with DiSC_3_(5) in a population assay. **A, E)** Representative fluorescence traces (AFU: arbitrary fluorescence units) of Em through time are shown. Each experiment displays its calibration curve. The addition of NH_4_Cl 10 mM activates CatSper channel causing depolarization by Ca^2+^ influx (left traces). **A)** Addition of 10 μM Secramine A previous to NH_4_Cl 10 mM led to a decrease in CatSper activation induced by alkalinization. **E)** Addition of 10 μM CASIN caused a marked depolarization by itself. In these experiments, calibration was not possible. **B, F)** Normalized △Em represents the difference between Em after NH_4_Cl addition (Em_NH4CI_) and before (resting: EmR or Em_Cdc42_ inhibitor) compared to the mean obtained in the control condition (without Cdc42 inhibitor addition). Data represents the mean ± SEM of at least 3 independent experiments. ΔEm of NC = 14.91 ± 0.96 mV (n = 11). ** p<0.01, * p<0.05 represents statistical significance vs. control (NC). Unpaired t-test was performed. **C, G)** Representative whole-cell currents traces from non-capacitated mouse sperm without addition of GTP. Inward and outward currents were elicited by a voltage ramp protocol from a holding potential of 0 mV (inset). To ensure stable recording conditions, first baseline currents (in HS solution) were obtained. Under HS condition (black traces) CatSper currents were minimal. In cation divalent-free (DVF) medium (dark gray traces) typical CatSper monovalent currents can be recorded. **C)** DVF after adding 10 mM NH_4_Cl, with (light gray traces) or without (blue traces) 10 μM Secramine. **G)** DVF after adding 10 mM NH_4_Cl, with (light gray traces) or without (blue traces) 10 μM CASIN. **D, H)** Quantification of *I_CatSper_* current densities. Secramine A (10 μM) or CASIN (10 μM) did inhibit the alkalinization-activated CatSper current at both negative and positive potentials (−80 and +80 mV). Data represents mean ± SEM of 4 sperm from different mice. ** p<0.01, * p<0.05 represents statistical significance vs. DVF control (+80 mV). ## p<0.01, # p<0.05 represents statistical significance vs. DVF control (−80 mV). One-way ANOVA was performed in combination with Holm-Sidak’s multiple comparisons test.

**Suppl. Figure 6:**
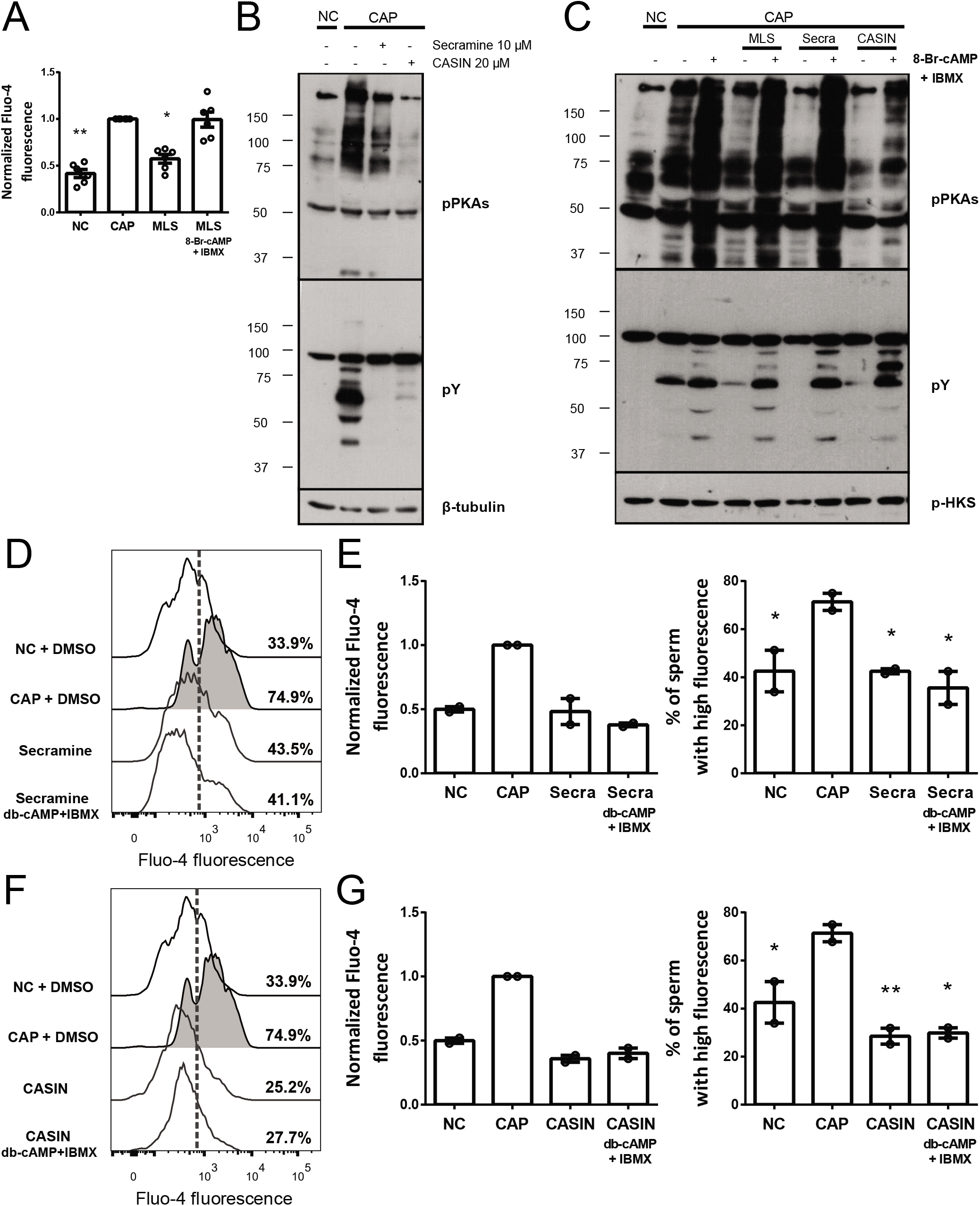
cAMP/PKA pathway inhibition was bypassed by using membrane permeable analogs of cAMP, but not the decrease in [Ca^2+^]_i_. **A)** Sperm incubated for 90 min under non-capacitating (NC) or capacitating conditions (CAP) in the absence (DMSO) or presence of 20 μM MLS-573151 were analyzed. 1 mM 8-Br-cAMP + 0.2 mM IBMX were used. Normalized MFI of Fluo-4 compared to the control condition (CAP 90 min with DMSO). Values represent the mean ± SEM of at least 6 independent experiments. **** p<0.0001, ** p<0.01, * p<0.05 represents statistical significance vs. control (CAP 90 min with DMSO). Non-parametric Kruskal-Wallis test was performed in combination with Dunn’s multiple comparisons test. **B-C)** Protein extracts were separated by 10% SDS-PAGE and immunoblotted with specific antibodies against phosphotyrosine residues (pY) and substrates phosphorylated by PKA (pPKAs). As a loading control, anti-β-tubulin or phosphor Hexokinase (p-HKS) (constitutively phosphorylated on tyrosine) were used. Representative images of at least 4 independent experiments are shown. Sperm were incubated for 90 min under non-capacitating (NC) and capacitating conditions (CAP) in the absence (DMSO) or presence of Cdc42 inhibitors (Secramine or CASIN). **B)** Incubation with 10 μM Secramine or 20 μM CASIN, resulted in a decrease in the pPKAs/pY phosphorylation patterns. **C)** Sperm were incubated under CAP conditions with 20 μM MLS-573151, 10 μM Secramine or 20 μM CASIN. The inhibition observed in this concentration was bypassed by using 1 mM 8-Br-cAMP + 0.2 mM IBMX. **D-G)** Ca^2+^ was measured in mouse sperm incubated for 90 min under non-capacitating (NC) or capacitating conditions (CAP) in the absence (DMSO) or presence of 10 μM Secramine or 20 μM CASIN. 1 mM db-cAMP + 0.2 mM IBMX were used. Representative histograms of normalized frequency vs. Fluo-4 fluorescence of non-PI stained sperm (live), with the corresponding percentage of sperm that increased Fluo-4 fluorescence, are shown (left panel) **(D, F)**. The percentage of sperm that responds by increasing the [Ca^2+^]_i_ was established in the CAP control condition and extrapolated to the others conditions (dashed line). Normalized MFI of Fluo-4 compared to the control condition (CAP 90 min with DMSO). Values represent the mean ± SEM of 2 independent experiments. Non-parametric Kruskal-Wallis test was performed in combination with Dunn’s multiple comparisons test. Percentage of sperm that increased Fluo-4 fluorescence after being incubated for 90 min in the different conditions. Values represent the mean ± SEM of 2 independent experiments. ** p<0.01, * p<0.05 represents statistical significance vs. control (CAP 90 min with DMSO). One-way ANOVA with Dunnett’s multiple comparisons test was performed.

**Suppl. Figure 7:**
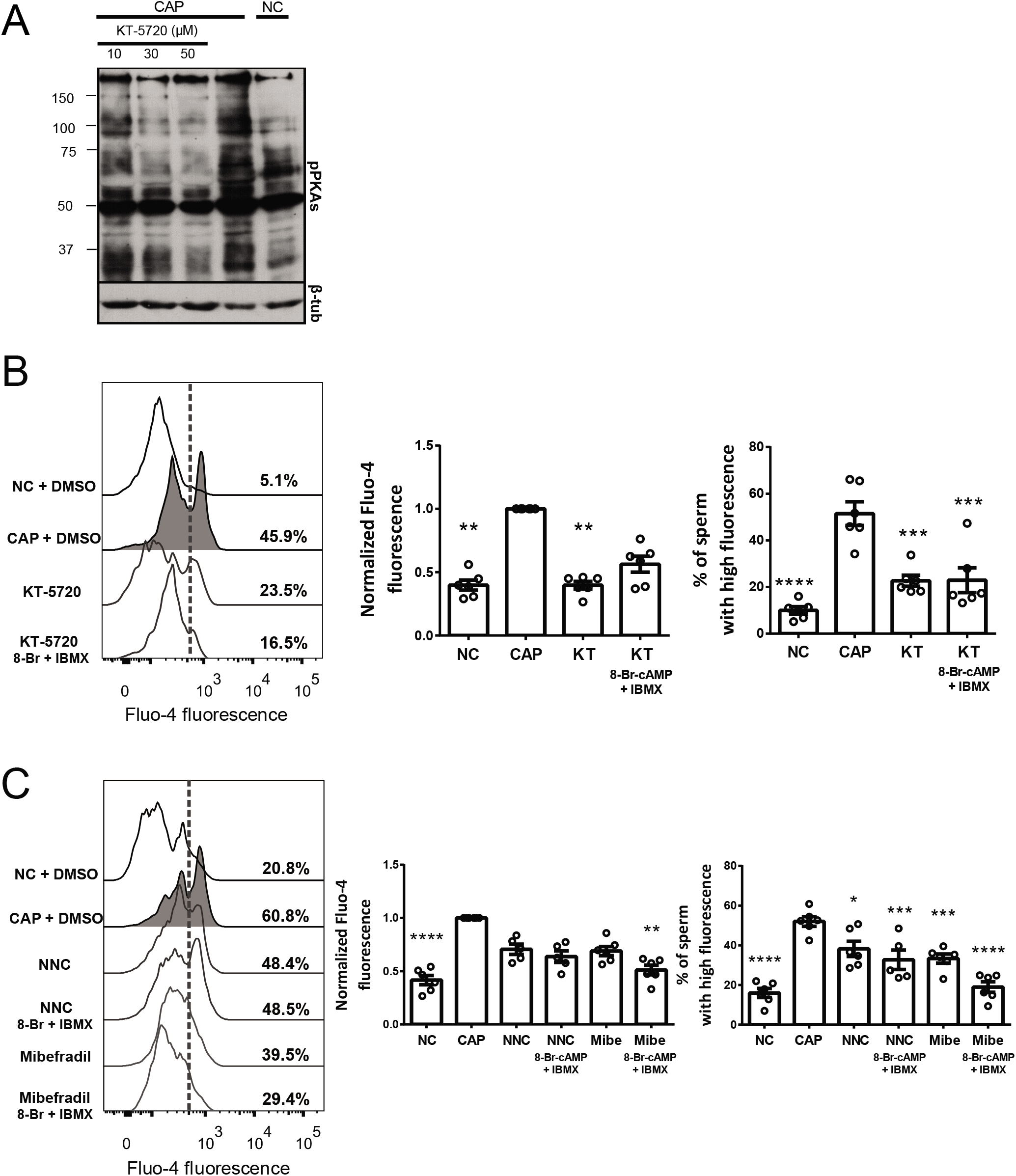
PKA and CatSper inhibition decreased [Ca^2+^]_i_. **A)** Protein extracts were separated by 10% SDS-PAGE and immunoblotted with antibodies against substrates phosphorylated by PKA (pPKAs). As a loading control, anti-β-tubulin was used. Increasing amounts of PKA inhibitor KT-5720 resulted in concentration-dependent decrease in pPKAs phosphorylation. Representative images of 3 independent experiments are shown. **B-C)** Ca^2+^ was measured in sperm incubated for 90 min under non-capacitating (NC) or capacitating conditions (CAP) in the absence (DMSO) or presence of the appropriate inhibitor: 30 μM KT-5720 (**B**), 10 μM NNC55-0396 or 30 μM Mibefradil (**C**). 1 mM 8-Br-cAMP + 0.2 mM IBMX were used. Representative histograms of normalized frequency vs. Fluo-4 fluorescence of non-PI stained sperm (live), with the corresponding percentage of sperm that increased Fluo-4 fluorescence, are shown (left panel). The percentage of sperm that responds by increasing the [Ca^2+^]_i_ was established in the CAP control condition and extrapolated to the others conditions (dashed line). Normalized MFI of Fluo-4 compared to the control condition (CAP 90 min with DMSO). Percentage of sperm that increased Fluo-4 fluorescence after being incubated for 90 min in the different conditions (right panel). Values represent the mean ± SEM of at least 5 independent experiments. **** p<0.0001, *** p<0.001, ** p<0.01, * p<0.05 represents statistical significance vs. control (CAP 90 min with DMSO). Non-parametric Kruskal-Wallis test was performed in combination with Dunn’s multiple comparisons test for MFI. One-way ANOVA with Dunnett’s multiple comparisons test was performed for data regarding percentage of sperm.

*Suppl. Video 1:* **Cdc42 is localized in four longitudinal lines.** 3D STORM reconstruction of Cdc42 in principal piece.

**Suppl. Table 1:**
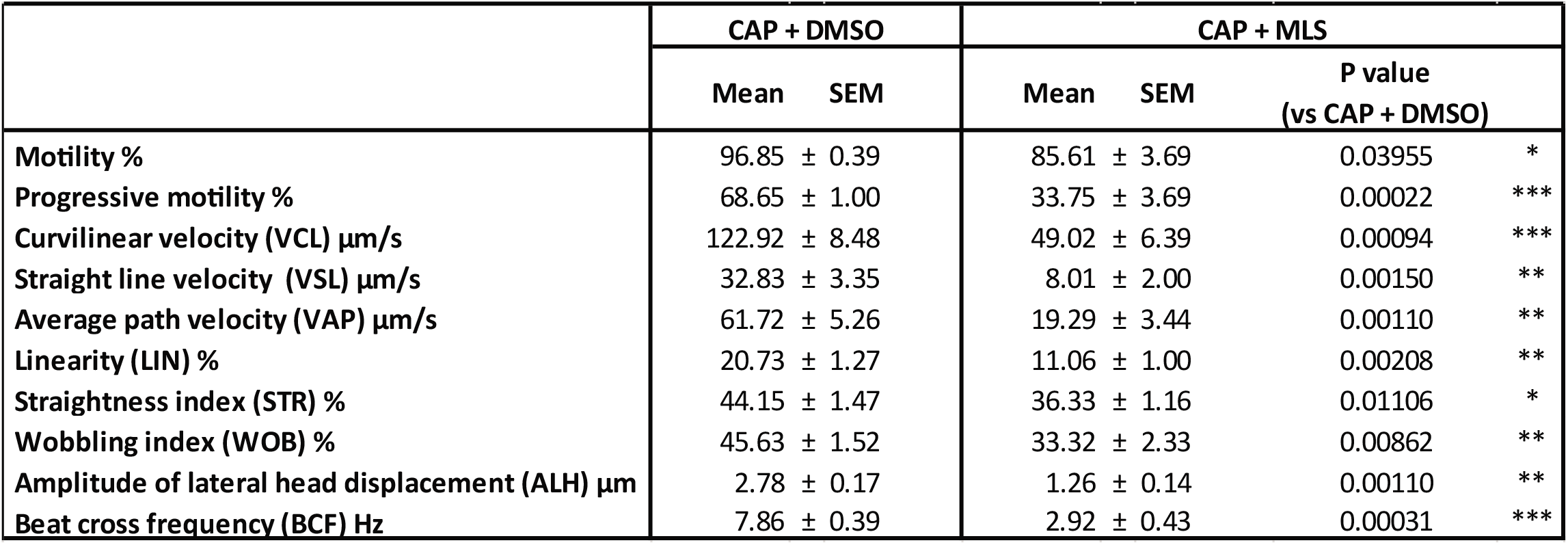
Effect of Cdc42 inhibition on sperm motility. CASA analysis was performed in sperm incubated during the 90 min of capacitation either in the absence (DMSO) or presence of 20 μM MLS-573151. Data represents the mean ± SEM of 4 independent experiments. *** p<0.001, ** p<0.01, * p<0.05 represents statistical significance vs. control (CAP 90 min with DMSO). Paired t-test was performed.

